# Opto-chemogenetic inhibition of L-type Ca_V_1 channels in neurons through a membrane-assisted molecular linkage

**DOI:** 10.1101/2024.05.12.593655

**Authors:** Jinli Geng, Yaxiong Yang, Boying Li, Zhen Yu, Shuang Qiu, Wen Zhang, Shixin Gao, Nan Liu, Yi Liu, Bo Wang, Yubo Fan, Chengfen Xing, Xiaodong Liu

## Abstract

Acute, specific, and robust inhibition of L-type Ca^2+^ (Ca_V_1) channels has been sought after for both research and therapeutic applications. Compared to other available Ca_V_1 antagonists, genetically-encoded modulators, such as CMI (C-terminus mediated inhibition) peptides encoded by Ca_V_1 DCT (distal C-terminus), hold great potentials due to its affirmative mechanisms of action on both gating and signaling. Here, we find that membrane-anchoring with a Ras tag could essentially help form a type of intramolecular-equivalent linkage, by which the tag anchored-peptide appears to dimerize with another protein or peptide on the membrane, supported by the evidence from patch-clamp electrophysiology and FRET imaging. We then design and implement the constitutive and inducible CMI modules, with appropriate dynamic ranges targeting the short and long variants of CaV1.3, both naturally occurring in neurons. Upon optical (infrared-responsive nanoparticles) and/or chemical (rapamycin) induction of FRB/FKBP binding, DCT peptides with no CMI in the cytosol acutely translocate onto the membrane via FRB-Ras, where the physical linkage requirement could be fulfilled. The peptides robustly produce acute and potent inhibitions on both recombinant CaV1.3 channels and neuronal CaV1 activities, and thus the Ca^2+^ influx-neuritogenesis coupling. Validated through opto-chemogenetic induction, this prototype demonstrates channel modulation via membrane-assisted molecular linkage, promising broad applicability to diverse membrane proteins.

## Introduction

L-type voltage-gated (CaV1.1-1.4) calcium channels are membrane proteins that mediate Ca^2+^ influx into excitable cells in response to transmembrane potentials ^1,2^. CaV1 channels play critical roles in a variety of pathophysiological processes related to muscle contraction, vesicle secretion, and gene transcription in cardiomyocytes or neurons ^3^. In particular, CaV1.3 and its close relative CaV1.2 are involved in the diseases (channelopathy) such as sinoatrial node dysfunction and deafness (SANDD) syndrome ^4,5^, adrenal hypertension ^6^, and Timonthy syndrome ^7,8^, all of which are due to the mutations on these channel genes. CaV1.3 and CaV1.2 are also implicated in neurodegenerative diseases including Parkinson’s disease and Alzheimer’s disease ^9,10^. Therefore, CaV1 agonists and antagonists have drawn significant interest and attention in both fundamental research and therapeutic development ^11–13^.

C-terminus mediated inhibition (CMI) is one emerging modality of CaV1 antagonism ^14–17^, which is closely related to calmodulin (CaM) modulation of the channels ^15^. Calmodulin, a Ca^2+^-binding protein containing EF-hands, is the molecular moderator of Ca^2+^-dependent inactivation (CDI); and its Ca^2+^-free form (apoCaM) is able to upregulate channel activation. Upon depolarization, the apoCaM-bound channel opens to induce Ca^2+^ influx and subsequently inactivates by conformational rearrangements associated with Ca^2+^-bound CaM (Ca^2+^/CaM). It has been proposed that the CaV1 channel is switched between the two states of high-versus low-opening controlled by apoCaM and Ca^2+^/CaM respectively ^14,18,19^. In this context, CMI sets the channel to its low-opening state, which is functionally and quantitatively equivalent to CDI or the low-opening state ^14^. In detail, two key motifs of distal C-terminus (DCT) in the pore-forming subunit (e.g.,α1D of CaV1.3), the proximal and distal C-terminal regulatory domains (PCRD and DCRD, respectively), compete against CaM by cooperatively binding the IQ domain (also known as CaM-binding domain, CaMBD) of the channel, where apoCaM is otherwise pre-associated ^14,19^. Such competitive binding underlying CMI serves as a unified principle applicable to all the DCT variants across the CaV1 family, only varying in their binding affinities ^17^. As one central principle of CMI, a physical linkage of any two modules among PCRD, DCRD and CaMBD (representing the channel with no DCT) is required and sufficient to induce CMI when the third module is present in the cell. The requirement of physical inter-module connection could be fulfilled either by constitutive linkage such as the fusion of PCRD with CaMBD or with DCRD, or by acute connection such as rapamycin-induced FRB/FKBP binding ^14^.

In this study, we aim to employ the above knowledge and tools to develop applicable CMI-based antagonists, for which high inhibition potency and ample dynamic range with acute induction are desired. However, such task is challenging in several aspects. Firstly, both short (without DCT) and long (with DCT) CaV1.3 variants are expressed in brain and heart cells ^20–24^. For the short variant of CaV1.3 (42A or α1DS), to achieve high CMI potency, it is necessary to utilize an engineered form of the channel, rather than its native form ^14^, to fulfill the requirement of “three-module principle” as mentioned above. Meanwhile, for the long form CaV1.3 (α1DL), the dynamic range is rather limited as DCT peptides directly result in either potent inhibition or no/minor CMI ^14,17^. The breakthrough began with our discovery of an unexpected, effective membrane-assisted connection between CMI modules. With the aid of the membrane-targeting CAAX tag from the Ras protein ^25^, aforementioned bottleneck problems could all be resolved by bringing the CMI modules onto the membrane. Lastly, for acute CMI induction, our previous rapamycin-inducible system or chemo-genetics in general provides a strong foundation for the development of the prototype ^14^, but with intrinsic limitations in spatiotemporal resolution, for which we proposed an approach by integrating with near infrared (NIR)-responsive nanoparticles, to release rapamycin and thus induce CMI.

In this work, we have developed a prototype of opto-chemogenetics, which consists of NIR-responsive rapamycin-encapsulated nanoparticles, and FRB/FKBP- and Ras-tagged peptides. Induction of membrane translocation helps form equivalent linkages to the targeted CaV1 channels, which potently inhibits CaV1 activities and neuronal development.

## Results

### Unexpected CMI on αDS by membrane-anchored P+D peptides

For the representative channel of short CaV1.3 (splice variant 42A, i.e.,α1DS) (Figure 1A, top), due to the lack of DCT, each channel (around IQ) is bound with apoCaM, thus supposedly having the maximum opening and ensuing inactivation ^14,18^, the latter of which was verified by examining the CDI strength of the whole-cell Ca^2+^ current (Figure 1A, middle). Briefly, the Ca^2+^ current (*ICa*, in red) of α1DS upon depolarization reached its instantaneous peak, and rapidly (within tens of milliseconds) inactivated until reaching its steady-state plateau. In contrast, nearly no or rather weak inactivation/decay was observed from the Ba^2+^ current (*IBa*, in gray) during the 300 ms depolarization step. The differences between Ca^2+^ currents and Ba^2+^ currents reflect the dependency of inactivation on Ca^2+^, quantified by the index of CDI or *SCa* (Figure 1A, bottom). According to the prerequisite of CMI, both PCRD and DCRD are required to be present for eventual association with the channel at the IQ domain as the trio complex; moreover, if there is no physical linkage among the three modules, no CMI would be produced ^14,17^. As expected, when PCRD and DCRD (encoding corresponding motifs of CaV1.3/1.4 DCT, denoted as P+D) were co-expressed with α1DS, the Ca^2+^ currents were indistinguishable from the control group (Figure 1B). Next, the Ras tag, as the membrane-targeting signal ^25^, was fused onto PCRD, DCRD or both, which were then co-expressed with the other untagged modules for further examination. Unexpectedly, when PCRD was anchored to the membrane (by CAAX motif of the Ras tag), CDI was significantly reduced (Scheme I, Figure 1C), compared to the control level of CDI obtained from α1DS alone (Figure 1A). Similar results of strong CMI effects on CDI were obtained with Ras-tagged DCRD (Scheme II, Figure 1D), and with co-expression of both Ras-tagged PCRD and Ras-tagged DCRD (Scheme III, Figure 1E). To quantify the inhibitory effects on α1DS, *CMI* (in percentage) is defined as the normalized fraction of channels that switch from apoCaM-bound to DCT-bound according to our previous study ^17^. *CMI* is inversely proportional to *SCa*, therefore substantial reduction of *SCa* corresponds to high *CMI*. All three schemes exhibited remarkable *CMI* (Figure 1F), apparently breaking the rule of CMI in that no bipartite linkage seemed to exist among the modules expressed in the cell. Meanwhile, except for the leakage requirement, the ‘trio’ working model of CMI is still valid. If only the PCRD or DCRD peptides were expressed alone, even if tagged with Ras—thereby lacking the third party—the channels did not exhibit any significant change in CDI (Figure S1). Taken together, these results regarding P+D peptides led us to hypothesize that the bipartite linkage requirement could be met by way of the Ras tag and the membrane, presumably as an unconventional type of ‘physical’ linkage allowing the CMI modules to form effective inter-module connections.

**Figure 1.**
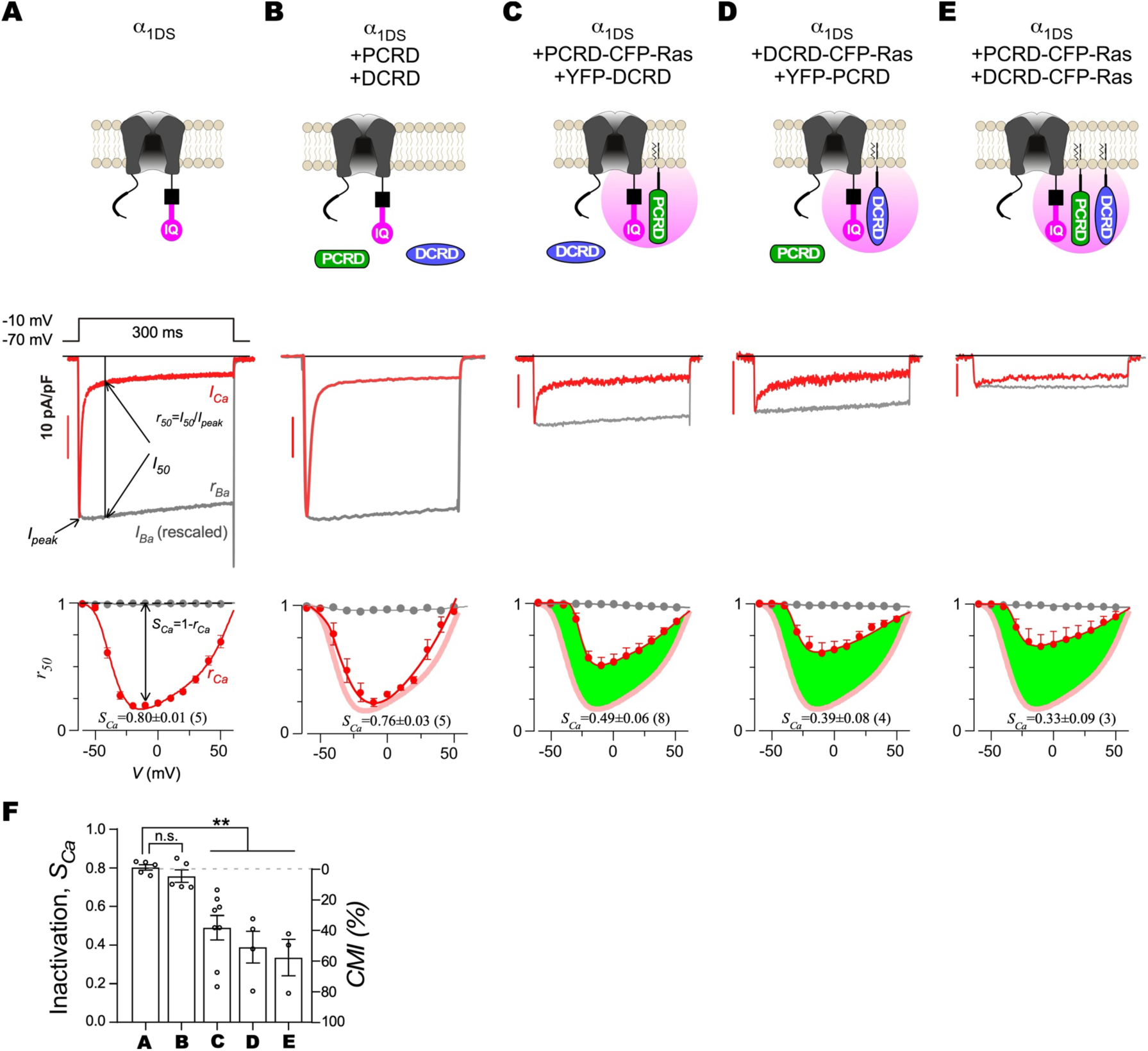
CMI effects on α1DS by P+D peptides. (A) Top: cartoon illustration of the CaV1.3 (splice variant 42A, denoted as α1DS) channel. Middle: representative current traces of α1DS at the membrane potential (*V*) of -10 mV. Ca^2+^ current (*ICa*, with the scale bar on the left) and Ba^2+^ current (*IBa*, rescaled to *ICa*) are colored in red and gray, respectively. The peak current (*Ipeak* representing the peak amplitude in pA or alternatively the current density in pA/pF) and the current at 50 ms (*I50*) are to calculate *r50* (ratio between *I50* and *Ipeak* for both *ICa* and *IBa*, i.e., *rCa* and *rBa*). Bottom: inactivation (*r50*) profiles across the full range of *V*. The differences between *rCa* and *rBa* reflect CDI (Ca^2+^-dependent inactivation), for which *SCa* (defined as 1-*rCa* at -10 mV) serves as the major index of CDI strength. (B) The negative control: two separate DCRD and PCRD (i.e., P+D) peptides in the cytoplasm. Both CFP-DCRD and YFP-PCRD were expressed together with α1DS channels. The schematic illustration, current traces and *r50* profiles are shown. The inactivation *rCa* profiles of the α1DS group (in pale red) versus the group of cytosolic P+D peptides (in red) are compared. (C) The design scheme (Scheme I) of PCRD on the membrane anchored by the Ras/CAAX tag while DCRD in the cytosol. The inactivation *rCa* profiles of the control group (in pale red) versus the peptide group (in red) are compared, for which their differences are highlighted by green shade. (D) The design scheme (Scheme II) of DCRD on the membrane while PCRD in the cytosol. (E) The design scheme (Scheme III) of anchoring both DCRD and PCRD on the membrane. (F) Statistical summary of CDI strength (*SCa*) and CMI potency (*CMI* in percentage). CMI potency is defined as the change in CDI: (*SCa,Control* - *SCa,Peptide*)/*SCa,Control*, where the control is α1DS. All statistical data are given as mean ± SEM (Standard Error of the Mean), with One-way ANOVA followed by Dunnett for post hoc tests: **, *p*<0.01; n.s., or not significant, *p*>0.05. See also Figure S1.

### Additional evidence from FRET supports the effective membrane-assisted linkage

Although electrophysiological recordings for P+D peptides have already strongly suggested that the membrane-targeting CAAX motif is critical to effectively link PCRD to DCRD or the channel, more direct evidence was sought after, for which Förster resonance energy transfer analysis (FRET) would serve as one suitable tool to address the molecular interactions in the cytosol or membrane ^26,27^ (Figure 2A-2C). Cytosolic peptides of P+D were co-expressed as a FRET pair, i.e., YFP-PCRD and CFP-DCRD, which resulted in low FRET efficiency (indexed by FRET ratio, *FR*), consistent with the weak *CMI* (Figure 1). By employing the membrane-targeting Ras tag, PCRD-YFP-Ras and DCRD-CFP-Ras peptides were constitutively expressed on the membrane (Figure 2A and 2B). Notably, Ras-tagged P+D peptides exhibited much higher *FR* than the cytosolic peptides (Figure 2C), indicating that a substantial number of donor and acceptor molecules are within the distance of 50Å according to the *r0* (Förster radius of FRET pair) between CFP and YFP ^28^. For the pair of membrane-anchored P+D that exhibited high *FR*, the PCRD and DCRD peptides are in such close proximity that they seem to be physically connected, apparently by a linkage equivalent to the one in the fusion protein of P-D. As expected, membrane-anchored CFP-YFP-Ras dimer resulted in high *FR*. Notably, after inserting an ER/K linker ^29^ into such dimer, i.e., CFP-ER/K-YFP-Ras, only produced low FRET thus serving as an important negative control (Figure 2D-2F). Assured by these control groups, intriguing results were obtained from CFP-Ras and YFP-Ras. This Ras-tagged FRET pair exhibited high *FR* on the membrane, mirroring the high *FR* observed from the PCRD-YFP-Ras and DCRD-CFP-Ras pair (Figure 2C). The similar results between Figure 2C and Figure 2F (indicated by the pink areas) argue no or ultraweak interaction present between PCRD and DCRD ^19^ consistent with our earlier reports (but see ^30^); otherwise, overall higher *FR* in Figure 2C should be expected because of additional PCRD/DCRD interactions. Combining electrophysiology and FRET results, we conclude that the P+D peptides with no cytosolic CMI, when anchored onto the membrane, effectively connect α1DS channels, thus fulfilling the physical linkage requirement for strong CMI on α1DS channels.

**Figure 2.**
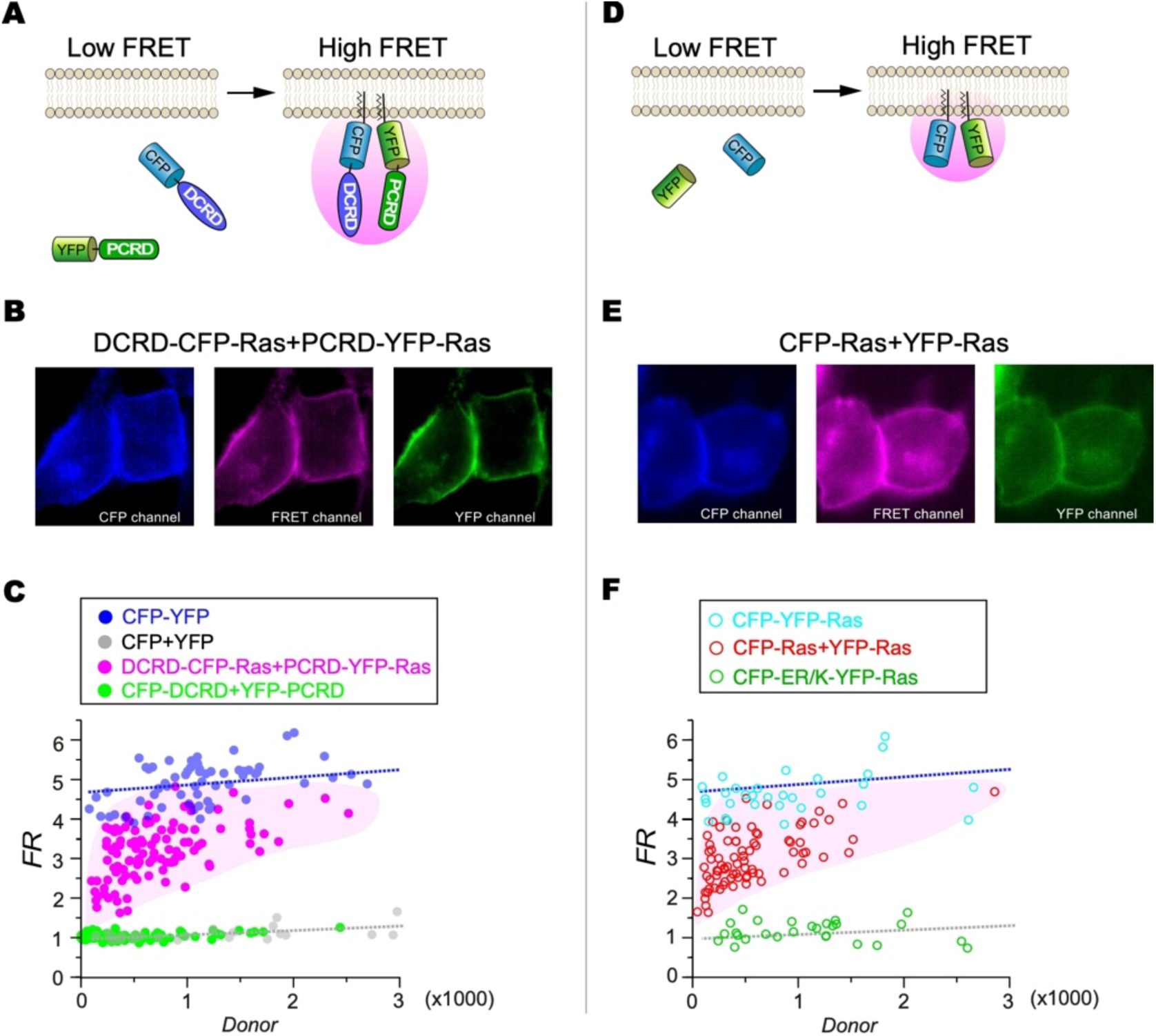
High FRET efficiency promoted by membrane-anchoring Ras/CAAX tag. (A) Schematic diagram of cytosolic and membrane-anchored FRET pairs of CFP-DCRD and YFP-PCRD. Once DCRD-CFP and PCRD-YFP are anchored onto the membrane by the Ras/CAAX tag, a putative molecular linkage (pink area) appears to be formed. (B) FRET imaging to examine the relationship between DCRD-CFP-Ras and PCRD-YFP-Ras. (C) Quantitative 2-hybrid 3-cube FRET describing the relationship between the FRET ratio (*FR*) and the donor fluorescence intensity (or concentration, *Donor*). Each dot indicates one HEK293 cell. The two FRET pairs of co-expressed P+D, i.e., Ras-tagged (pink) versus cytosolic (green) CFP-DCRD and YFP-PCRD, were compared. The CFP-YFP dimer and CFP/YFP co-expression serve as the positive (blue) and negative (gray) control, respectively. (D) Schematic diagram of cytosolic and membrane-anchored FRET pairs of CFP and YFP. The pink area indicates potential linkage between CFP and YFP. (E) FRET imaging to examine the relationship between CFP-Ras and YFP-Ras. (F) The FRET pairs were examined including CFP-Ras and YFP-Ras (co-expression, red), CFP-YFP-Ras (dimer, cyan), CFP-ER/K-Ras (dimer, dark green). The two control lines (gray and blue) are directly adopted from (C).

### Design of P-D peptides for α1DL inspired by membrane-assisted CMI on α1DS

So now we have partially achieved our goal by providing P+D peptides which, with the assistance of the membrane, target α1DS with high potency. With respect to α1DL, which is the full-length variant of CaV1.3, the potency of CMI is expected to correlate quantitatively with the affinity between DCT peptides and α1DL channels. Accordingly, in contrast to the DCT peptides encoded by CaV1.3 or CaV1.4, the DCT peptides encoded by CaV1.1- or CaV1.2 such as CCATC containing the linked PCRDC and DCRDC of CaV1.2 are barely able to produce any inhibition on α1DL ^17^. This type of peptide is denoted as P-D (to compare with P+D), since PCRD and DCRD are physically linked together. Consistent results were achieved that CDI of the CCATC group is indistinguishable from the α1DL control (Figure 3A and 3B). Inspired by the discovery that membrane-assisted molecular linkage significantly enhances CMI, we then proposed the design of Ras-mRuby-CCATC, where the Ras is fused, as a novel type of applicable P-D peptides. In contrast to cytosolic CCATC (or CaV1.2-encoded P-D), membrane-anchored CCATC significantly attenuated CDI of α1DL (Figure 3C), resulting in potent *CMI* of 57 ± 9% (Figure 3D). A major feature of CMI effects is the concurrent inhibition of both inactivation and activation processes. In support of this notion, membrane-anchored P-D exhibited smaller Ca^2+^ currents (pA/pF) in comparison with cytosolic P-D producing nearly no effect on α1DL (Figure S2). Thus, we have designed and implemented the CaV1.2-encoded P-D peptides that are highly applicable to α1DL. They produce nearly no inhibition in the cytosol, yet exhibit potent CMI effects when translocated onto the membrane.

**Figure 3.**
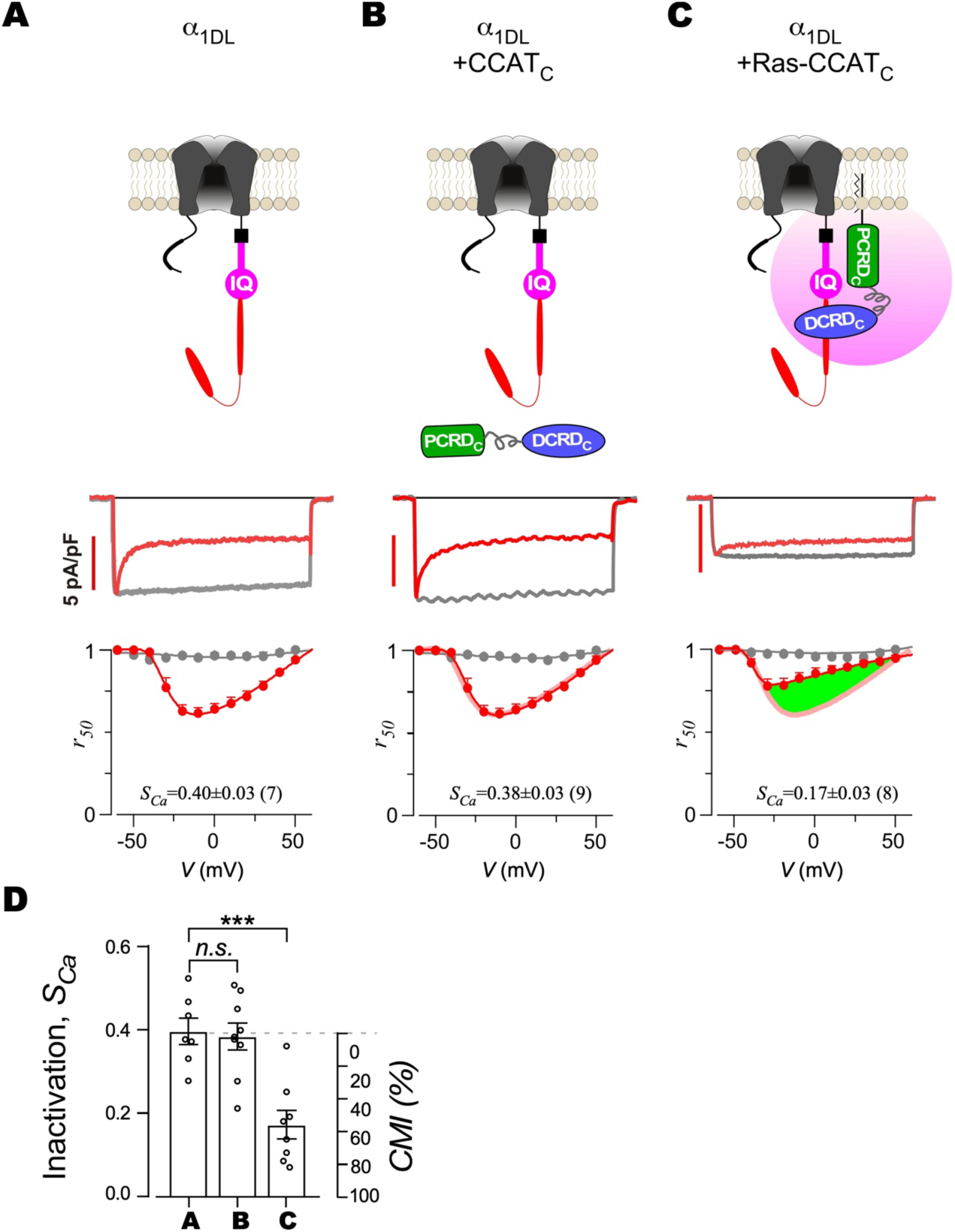
CMI effects of CaV1.2-encoded peptides (P-D) on α1DL. (A) Top: illustration of CaV1.3 long (denoted as α1DL). Middle: representative *ICa* and *IBa* traces of α1DL at the membrane potential (*V*) of -10 mV. Bottom: voltage-dependent inactivation profiles of *r50*. (B) The design scheme for the P-D type of peptide CCATC (P-D) encoded by CaV1.2 DCT. The inactivation *rCa* profiles of the α1DL group (in pale red) versus the group of cytosolic P-D peptides (in red) are compared. (C) The design scheme of membrane-anchored P-D peptides. α1DL channels were co-expressed with Ras-mRuby-CCATC. The inactivation *rCa* profiles of the group of P-D peptides (in red) versus the α1DL control group (in pale red) are compared, for which their differences are highlighted by green shade. (D) Statistical summary of the strength of CDI (*SCa*) and CMI potency (*CMI*). Data points are presented as mean ± SEM, with the corresponding significance: ***, *p*<0.001; n.s., *p*>0.05. See also Figure S2.

### Chemical and optical induction of rapid cytosol-membrane peptide translocation

In the context of membrane-assisted physical linkage for CMI, the induction method is crucial for CMI peptides to acutely translocate onto the membrane. Following the previous design of acute CMI ^14^, a series of chemical/rapamycin-inducible versions targeting α1DS were developed by introducing rapamycin binding peptides of FRB/FKBP alongside the membrane-anchoring Ras tag. The primary rationale behind such a design is to swiftly fulfill the requirement of physical linkage to the peptides (Figure 4A). By design, various DCT-encoded motifs in the cytosol are able to acutely translocate onto the membrane through the tight binding of rapamycin to FKBP-tagged peptides which would eventually form the complex with FRB-Ras on the membrane ^14,31^. YFP-FKBP-DCRD, YFP-FKBP-PCRD, and FRB-CFP-Ras were expressed in HEK293 cells. Time-lapse imaging by confocal microscopy was conducted to monitor the process of membrane translocation at the interval of 30 s (Figure 4B). To quantify the membrane translocation, a red line was drawn to cross the whole cell from which the ratio of fluorescence intensity between membrane (Fm) versus cytosol (Fc) was calculated as a quantitative index. Direct application of 1 μM rapamycin to the bath induces the binding of FRB fused in the membrane by Ras tag and FKBP, leading to translocation of YFP-FKBP-PCRD and YFP-FKBP-DCRD on the membrane, while treatment with 0.1% DMSO, serving as a vehicle control, did not cause any membrane association with FKBP (Figure 4B and 4C).

**Figure 4.**
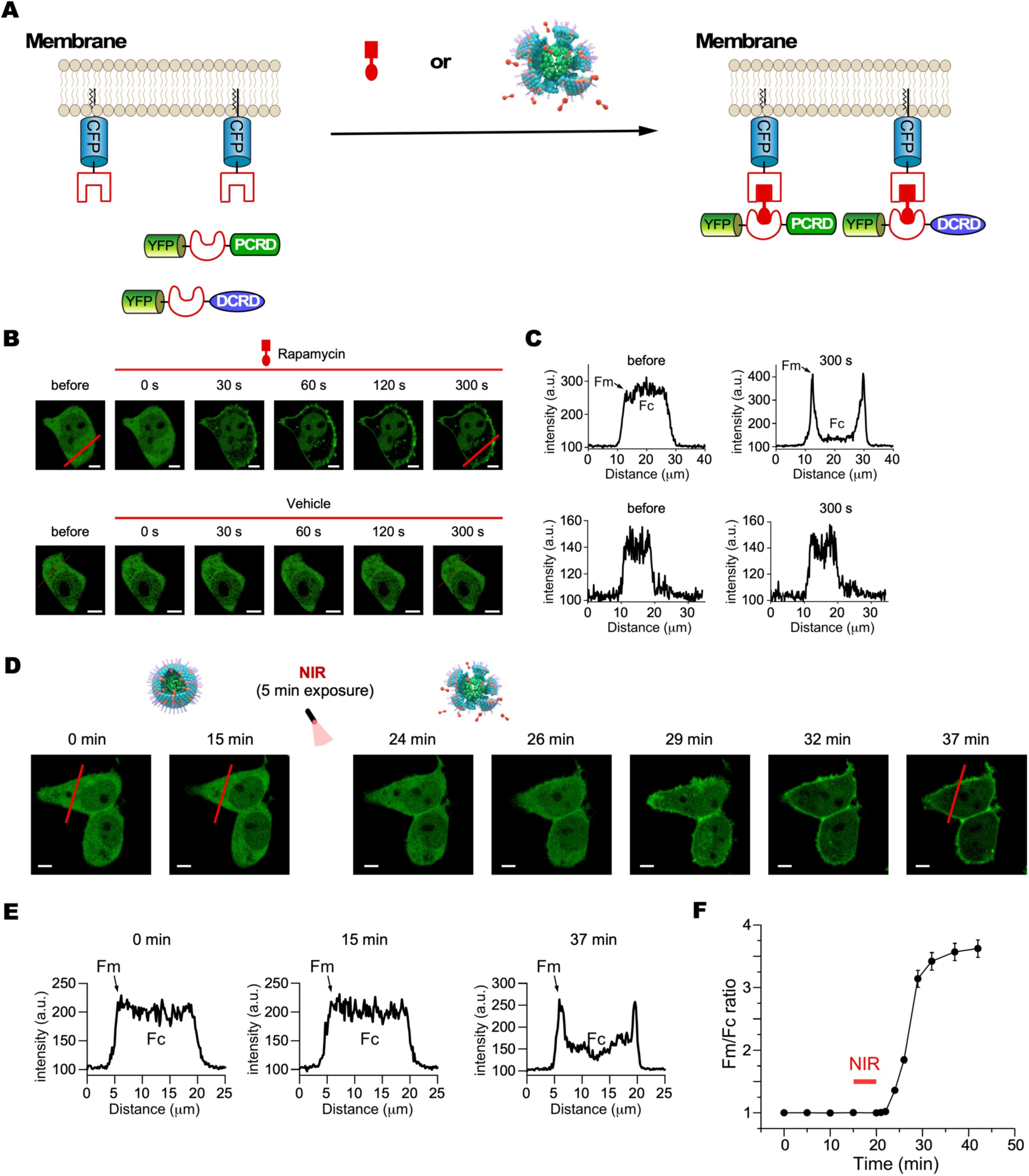
Chemical and optical induction of rapid cytosol-membrane translocation. (A) The general design scheme for chemical or optical induction of membrane translocation. Rapamycin or encapsulated rapamycin releasing from rapamycin@PDPP nanoparticles after 808 nm NIR laser irradiation induces FRB/FKBP heterodimerization. In response to chemical or optical induction, YFP-FKBP-DCRD and YFP-FKBP-PCRD (P+D) rapidly translocate from cytosol to the plasma membrane, where FRB-CFP-Ras is constitutively anchored on the membrane. (B) Representative confocal images to illustrate the rapid translocations of cytosolic DCRD and PCRD onto the plasma membrane within 5 min upon rapamycin induction. HEK293 cells were transfected with YFP-FKBP-DCRD, YFP-FKBP-PCRD, and FRB-CFP-Ras. 1 μM rapamycin or vehicle (0.1% DMSO, negative control) was applied. The scale bar is 5 μm. (C) The representative fluorescence intensity profiles before or 300 s after induction, based on the cross sections (red lines) in the images. Fm and Fc represent membranal fluorescence and cytosolic fluorescence, respectively. (D) Representative confocal images to illustrate NIR induction of translocations of cytosolic YFP-FKBP-DCRD and YFP-FKBP-PCRD (P+D) to the plasma membrane in real-time and *in situ*. 25 μg/ml rapamycin@PDPP nanoparticles were incubated with cells for 15 min and irradiated by NIR for 5 min to release the rapamycin reaching the final concentration of 1 μM *in situ*, then imaged for 22 min to monitor peptide translocations. The concentration of rapamycin@PDPP nanoparticles was calibrated according to the UV standard curve of conjugated polymer PDPP. The scale bar is 5 μm. (E) The representative fluorescence intensity profiles at 0 min, 15 min and 37 min. (F) Temporal profiles of the ratio of membranal fluorescence to cytosolic fluorescence (Fm/Fc). The red bar indicates 5 min of NIR exposure. All statistical data are given as mean ± SEM. See also Figure S3.

Our strategy for achieving optical induction fully leverages the previously established method. NIR-responsive nanoparticles encapsulating rapamycin (rapamycin@PDPP) may provide additional benefits in spatial control, tissue penetration, biocompatibility and drug release.

As shown in Figure S3, rapamycin@PDPP nanoparticles were prepared via the nanoprecipitation of PDPP, DPPC, DSPE-PEG2000-NH2 and rapamycin. PDPP cores convert NIR light into heat, triggering a phase-transition of DPPC lipid coating at a temperature of 41°C. An additional lipid coating, DSPE-PEG2000-NH2, enhances the biocompatibility of the nanoparticles. Encapsulated rapamycin is released from the nanoparticles under 808nm NIR irradiation. The NIR-responsive nanoparticles have been proven to be biocompatible in HEK293 cells and tissues ^32,33^. To approach the design goals, rapamycin@PDPP nanoparticles were then deployed to gain opto-chemical control of membrane translocation of CMI modules. The first key step was to develop NIR-responsive binding between FRB-Ras and FKBP. Fluorescent tag YFP was fused onto FKBP to visualize its subcellular distribution. As illustrated in the design scheme (Figure 4A), NIR light triggers the release of rapamycin from rapamycin@PDPP nanoparticles, and subsequently induces the binding (heterodimerization) of Ras-tagged FRB and cytosolic FKBP in the cell, leading to translocation of YFP-FKBP-PCRD and/or YFP-FKBP-DCRD onto the membrane. The concentration of rapamycin@PDPP nanoparticles was calibrated according to the UV standard curve of conjugated polymer PDPP. Considering the efficacies of drug loading and release of nanoparticles, cells were incubated with 25 μg/ml rapamycin@PDPP and irradiated by NIR for 5 min to release the rapamycin resulting in a final concentration of 1 µM rapamycin in the bath solution. No basal leakage of rapamycin from rapamycin@PDPP was detected without NIR stimulation during the 15-minute imaging period. After 5 minutes of *in situ* NIR exposure to trigger rapamycin release, we observed rapid mobilization of the YFP-tagged proteins (Figure 4D). As demonstrated by histogram analyses of positions along the red line, distinct intensity profiles were observed at 0 or 15 minutes (prior to NIR exposure) compared to 37 minutes (following NIR exposure) (Figure 4E). This temporal analysis of the Fm/Fc ratio further underscored the stability of rapamycin encapsulated within PDPP in the absence of NIR stimulation, and its rapid membrane translocation triggered by opto-chemical activation of FRB/FKBP dimerization (Figure 4F).

### Chemically and optically inducible membrane-assisted CMI peptides of P+D

Leveraging the membrane-assisted peptide linkage (Figure 1) and rapid induction of cytosol-membrane translocation (Figure 4), we designed a prototype of membrane-assisted CaV1.3-encoded P+D peptides, enabling CMI on α1DS (Figure 5A). Following our design, we used rapamycin or rapamycin@PDPP to acutely induce the translocation of YFP-FKBP-PCRD or YFP-FKBP-DCRD onto the membrane via Ras-tagged FRB. As anticipated, Ca^2+^ currents of α1DS were potently inhibited upon 1 μM rapamycin, demonstrated by the time-dependent inactivation (*SCa*) and activation (indexed by the current amplitude, *Ipeak*) (Figure 5B and 5C). In control α1DS channels lacking P+D expression, 1 μM rapamycin had no effect, as shown by unchanged *SCa* and *Ipeak* values (Figure S4A-S4C). Vehicle control (0.1% DMSO) did not impact Ca^2+^ currents when α1DS was co-expressed with inducible P+D peptides (Figure S4D-S4F).

**Figure 5.**
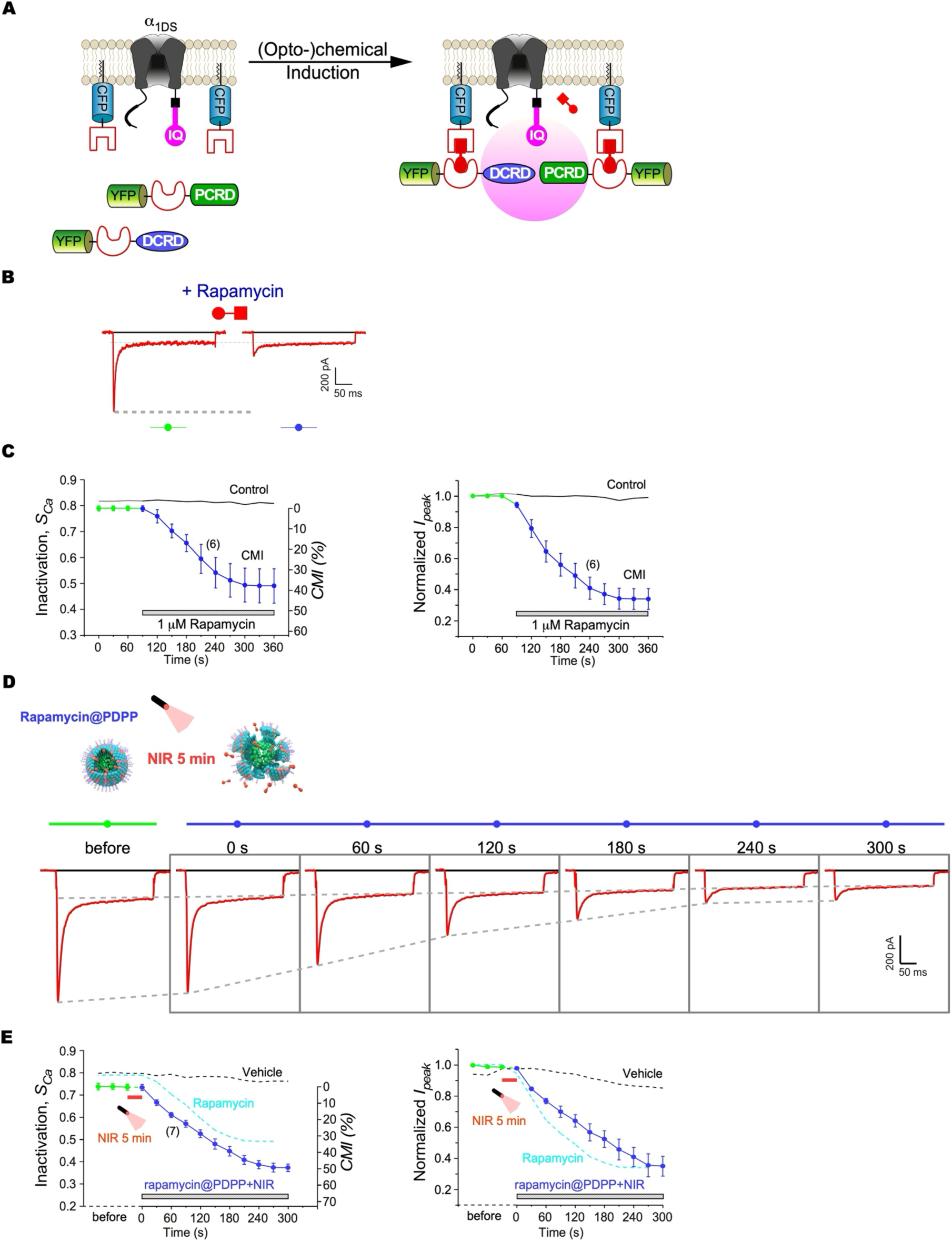
Opto-chemogenetics of membrane-assisted CMI on α1DS by P+D peptides. (A) The design scheme of chemogenetic or opto-chemogenetic CMI. Rapamycin or opto-chemical control of rapamycin releasing from rapamycin@PDPP nanoparticles is intended to induce rapid translocation of YFP-FKBP-DCRD and YFP-FKBP-PCRD from the cytosol to the plasma membrane. (B, C) Rapamycin-induced CMI through translocation of P+D peptides onto the membrane. YFP-FKBP-DCRD and YFP-FKBP-PCRD were co-expressed with FRB-CFP-Ras. Exemplars of Ca^2+^ current traces demonstrate effective inhibition by 1 μM rapamycin, where both the peak (*Ipeak*) and steady-state amplitude (at 300 ms, i.e., *I300*) are indicted by the dashed lines (B). Temporal profiles of *SCa* along with *CMI* (left) and normalized *Ipeak* (right) demonstrate rapamycin-induced attenuation in comparison with α1DS control (C). Data points connected by lines (in green and blue) represent the values of *SCa* or *Ipeak* before and after rapamycin application, respectively. (D) Opto-chemogenetic CMI for α1DS. HEK293 cells expressing recombinant α1DS, FRB-CFP-Ras, YFP-FKBP-PCRD, and YFP-FKBP-DCRD were treated with rapamycin@PDPP nanoparticles. NIR irradiation for 5 min to trigger the release of rapamycin reaching the final concentration of 1 μM. Representative Ca^2+^ traces at different timepoints are featured with a declining trend of *Ipeak* and the characteristic stable *I300*. (E) Temporal profiles of inactivation *SCa* along with potency *CMI* (left) and normalized *Ipeak* (right). Before (green) and after 5-min NIR exposure (blue), *ICa* currents were recorded every 30 s. Dotted lines in black and cyan represent the control groups of the vehicle (0.1% DMSO, details in Figure S4D-S4F) or rapamycin (direct application, adapted from Figure 5C). All statistical data are given as mean ± SEM. See also Figure S4-S7.

In principle, three possible schemes can be deployed according to the particular peptides translocated onto the membrane: PCRD (Scheme I), DCRD (Scheme II) or both (Scheme III). According to the first scheme (Figure S5A-S5C), the two DCT modules of FKBP-PCRD and DCRD were co-expressed with the third module, IQ-containing membrane channels. Upon administration of rapamycin, both inactivation and activation as indexed by *SCa* and *Ipeak*, were gradually attenuated, causing clear CMI effects (potency 14 ± 1%). Presumably, an effective connection formed between the channel and FKBP-PCRD, the latter of which was rapidly mobilized onto the membrane through rapamycin-triggered FKBP binding to membranous FRB-Ras. In the presence of cytosolic DCRD, the effectively-connected IQ (channel) and PCRD would further form a ternary complex according to the established principle of CMI. Consistently, despite the apparent contradiction of concurrent attenuation on both activation and inactivation, the net reduction of Ca^2+^ influx was ensured by the stereotypical *I300* (current amplitude at 300 ms), which remained unaltered (Figure S5A).

In the second scheme, similar rapamycin-induced CMI effects were observed from Ca^2+^ current recordings (Figure S5D-S5F), with FKBP-DCRD and cytosolic PCRD co-expressed alongside α1DS. Consistent with the data and interpretation from the first scheme, effective connections between FKBP-tagged DCRD and α1DS presumably formed in response to rapamycin, leading to cooperative binding and channel inhibition.

Notably, in the third scheme (Figure 5B and 5C), both PCRD and DCRD were recruited to the membrane, directly corresponding to its constitutive version (Figure 1E). The CMI potency of this scheme appeared relatively more pronounced than the other schemes. This was evident from the exemplar Ca^2+^ current traces before and after rapamycin stimuli, and from the temporal profiles of inactivation (*SCa*) and activation (*Ipeak*).

We then explored the optical induction of CMI, following our strategy of NIR-responsive rapamycin release and subsequent peptide translocation onto the membrane. Rapamycin@PDPP nanoparticles without NIR light did not cause any significant effect, indicated by unaltered CDI (*SCa*) and peak current (*Ipeak*) during the whole time-course (Figure S6A and S6B). Five minutes of NIR irradiation successfully induced inhibitory effects on Ca^2+^ currents (Figure 5D), presumably by triggering the release of rapamycin, leading to typical attenuation on both inactivation (*SCa*) and activation (*Ipeak*) (Figure 5E). Moreover, the CMI potency associated with optical induction here was close to the potency of CMI by direct rapamycin induction (cyan lines based on Figure 5B and 5C), further confirming the effectiveness of the combined optical/chemical induction as proposed in our design. As a negative control, rapamycin@PDPP nanoparticles with NIR light had no significant effect on HEK293 cells only expressing α1DS (Figure S6C and S6D).

Lastly, detailed profiles of *SCa* and *Ipeak* across the full range of membrane potentials were measured for Ca^2+^ currents toward the protocol’s end (Figure S7), yielding consistent conclusions in all cases, including the three basic schemes and the optical induction design (P+D) (Figure 5; Figure S5). The potencies for the Scheme I, II and III are 19 ± 2%, 36 ± 15% and 67 ± 9%, respectively, in addition to opto-chemogenetic CMI of 56 ± 9%.

### P-D peptides inducibly suppressed the CaV1 activity-neuritogenesis coupling

According to our newly designed P-D peptides (Figure 3), the CaV1.2-encoded P-D peptides of PCRDC-DCRDC (PCDC) on the membrane are able to produce CMI effects. An inducible version of P-D was developed using the rapamycin/FRB/FKBP module similar to what is depicted in Figure 5A. First, FKBP-YFP-PCDC and FRB-CFP-Ras were tested for the inducible CMI effects on α1DL. PCDC was recruited by acute induction to the membrane resembling its constitutive version (Figure 6A-6C). Upon rapamycin, both inactivation and activation as indexed by *SCa* (Figure 6B) and *Ipeak* (Figure 6C; Figure S8A) were gradually attenuated, causing unambiguous CMI effects (*CMI* 44 ± 7%), further confirmed by the full voltage-dependent profiles (Figure S8B). Interestingly, FKBP-YFP-PCDC also significantly induced strong CMI inhibition (41 ± 6%) on α1DS, evidenced by gradual attenuations in inactivation (*SCa*) and activation (*Ipeak*) (Figure 6D-6F). Toward the ending point of the time course, attenuated CDI of α1DS was close to that of α1DL (Figure 6A-6C). The affinity of PCDC for the channel should be enhanced by about two orders of magnitude before and after membrane-translocation ^17^, which could be well accounted for by a membrane-assisted dimerization-like linkage as proposed earlier.

**Figure 6.**
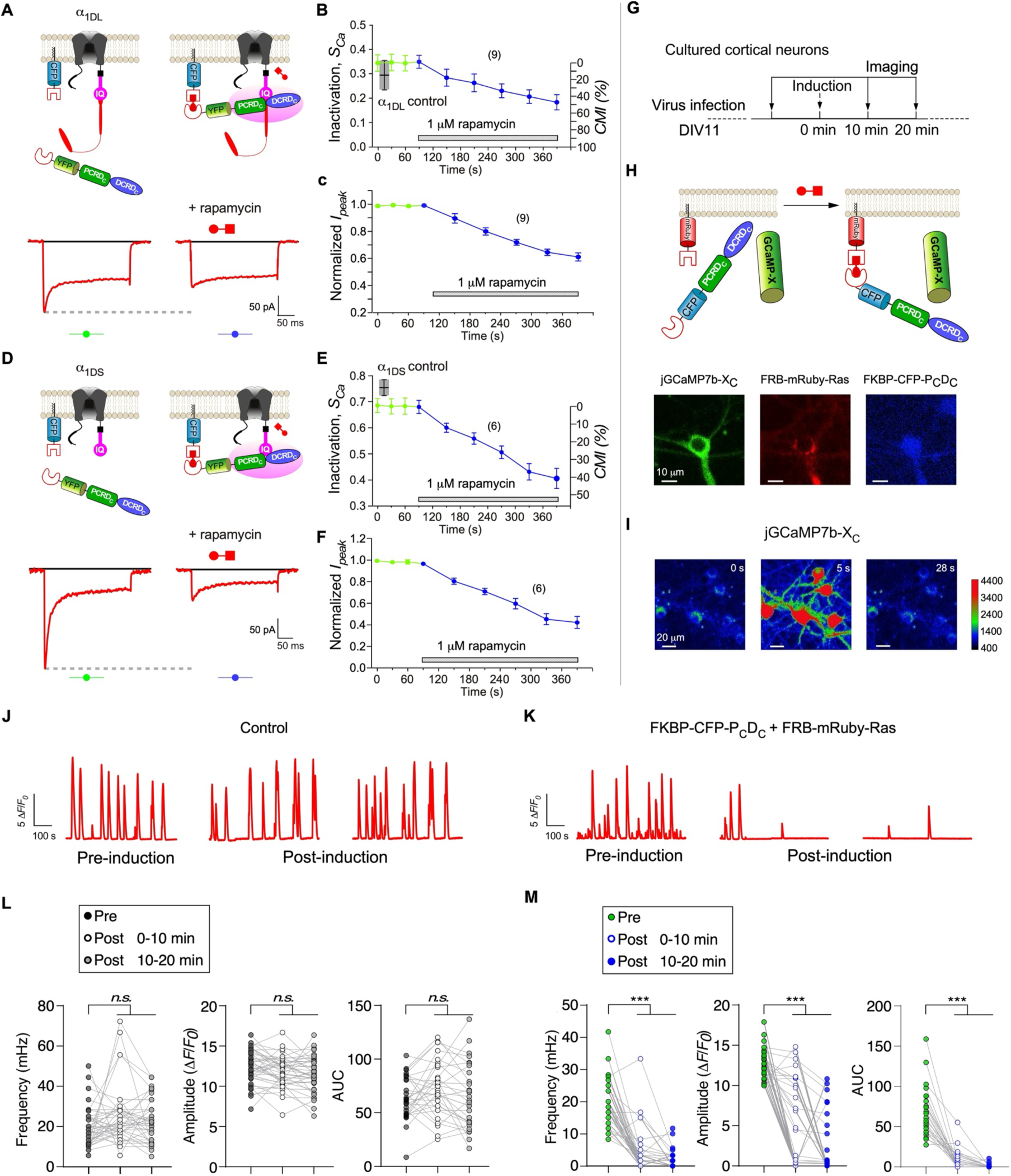
P-D peptides inducibly suppress recombinant CaV1.3 channels and neuronal Ca^2+^ oscillations. (A-C) Rapamycin-induced membrane-targeting of CaV1.2 DCT-encoded PCRDC-DCRDC (PCDC) and effective CMI for CaV1.3 long (α1DL). Membrane-anchored FRB-CFP-Ras recruited FKBP-YFP-PCDC onto the membrane by rapamycin-mediated FRB/FKBP binding (A). CMI was induced presumably by effective linkage between membrane-anchored PCDC and α1DL. Exemplars of Ca^2+^ current traces demonstrate the reduction in the peak amplitude (*Ipeak*, dotted line) by 1 μM rapamycin (A). Temporal files of *SCa* along with CMI potency (*CMI*) (B) and normalized *Ipeak* (C) are to show rapamycin-induced attenuation. Data points in green and blue (connected by lines) represent the values of *SCa* or *Ipeak* before and after rapamycin administration, respectively. (D-F) Rapamycin-induced membrane-targeting of PCDC and effective CMI for α1DS. Membrane-anchored FRB-CFP-Ras recruited FKBP-YFP-PCDC to the membrane by rapamycin-mediated FRB/FKBP binding (D). Effective CMI was induced presumably by effective connection between membranous PCDC and α1DS. Exemplars of Ca^2+^ current traces demonstrate the effective inhibition on *Ipeak* by 1 μM rapamycin (D). Temporal files of *SCa* and *CMI* (E) and normalized *Ipeak* (F) are shown, where data points in green and blue (connected by lines) represent the values of *SCa* or *Ipeak* before and after rapamycin application, respectively. (G) Experimental protocol for acute inhibition on cultured cortical neurons. 10 μM rapamycin was added to neurons expressing membrane-anchored FRB-mRuby-Ras and cytosolic FKBP-CFP-PCDC by lentivirus transfection at DIV 11. Confocal imaging was performed before and after rapamycin treatment at the indicated timepoints. (H) The Ca^2+^ probes, P-D peptides and induction modules were co-expressed in neurons, subject to confocal fluorescence imaging: jGCaMP7b-XC (green), FRB-mRuby-Ras (red), and FKBP-CFP-PCDC (blue). (I) Time-lapse confocal images of Ca^2+^ dynamics in cortical neurons at DIV 11. Color-coded fluorescence intensities (as shown in the scale bar) of jGCaMP7b-XC reflect the concentrations of cellular Ca^2+^ signals. (J, K) Spontaneous Ca^2+^ activities in a control neuron (J) or a neuron with rapamycin-inducible CMI (K). (L, M) Key indices to quantify Ca^2+^ dynamics in the control group (L) and the neurons expressing CMI modules (M), including the average frequency (mHz) (left), the peak amplitude (*ΔF*/*F0*) (middle), and the Ca^2+^ influx AUC (calculated as *ΔF*/*F0* per min) (right). All statistical data are given as mean ± SEM. One-way ANOVA followed by Dunnett for post hoc test is used for L, M: ***, *p*<0.001; n.s., *p*>0.05. See also Figure S8 and S9.

Our next step was to investigate inducible membrane-assisted P-D peptides in live neurons under physiological conditions, following the validation of CMI impacts on recombinant CaV1.3 channels. Furthermore, peptides encoded by DCT, including CCATC, are endogenously present in native cells, playing pivotal roles in the regulation of channels and neurons. Subsequently, FKBP-CFP-PCDC and FRB-mRuby-Ras were virally introduced into cultured cortical neurons. An improved version of GCaMP, jGCaMP7b-XC ^34^, was utilized to monitor neuronal Ca^2+^ dynamics. Following rapamycin induction, PCDC translocated onto the cortical neuron membrane (Figure 6G-6I). In cortical neurons expressing FKBP-CFP-PCDC and FRB-mRuby-Ras, we observed spontaneous Ca^2+^ oscillations, quantified using the indices of frequency (mHz), amplitude (Δ*F/F0*) and influx (AUC, or Area Under the Curve). Oscillatory Ca^2+^ activities post 10 and 20 minutes of rapamycin treatment were compared to pre-treatment levels (Figure 6J-6M). In control neurons not expressing FKBP-CFP-PCDC, no perceptible difference was observed (Figure 6J and 6I). Conversely, the membrane-anchored P-D type peptides of PCDC significantly reduced Ca^2+^ oscillations, as indicated by all three major indices (Figure 6K and 6M). Following rapamycin induction, hippocampal neurons also exhibited a similar reduction in spontaneous Ca^2+^ activities (Figure S9).

Neurite outgrowth and neuronal development are closely related to Ca^2+^, particularly due to CaV1 channels ^17,35^. To explore the CaV1 activity-neuritogenesis coupling in the context of inducible CMI, we examined cultured cortical neurons at different timepoints (days *in vitro*, DIV) after rapamycin treatment (Figure 7A and 7B). As expected, both neuritogenesis (total length) and Ca^2+^ influx (AUC) of cortical neurons expressing FKBP-CFP-PCDC and FRB-mRuby-Ras were significantly attenuated compared to the control group not expressing the CMI modules. Consistent with previous results ^34^, the neurite length of each neuron in accordance to its DIV followed a sigmoidal-like curve, as evidenced from the control (Figure 7C). We qualitatively assumed that CaV1 influx to be the key factor responsible for neuritogenesis, despite the complexity of frequency, amplitude and other parameters of Ca^2+^ dynamics ^34^. In this study, Ca^2+^ influx was explicitly quantified as AUC, represented as a bell-shaped curve (Figure 7D), aligning well with the two-phase trend of neurite growth rate (NGR, μm/day) (Figure 7E), thereby directly supporting a clear yet simple correlation between Ca^2+^ influx and neurite outgrowth (Figure 7F).

**Figure 7.**
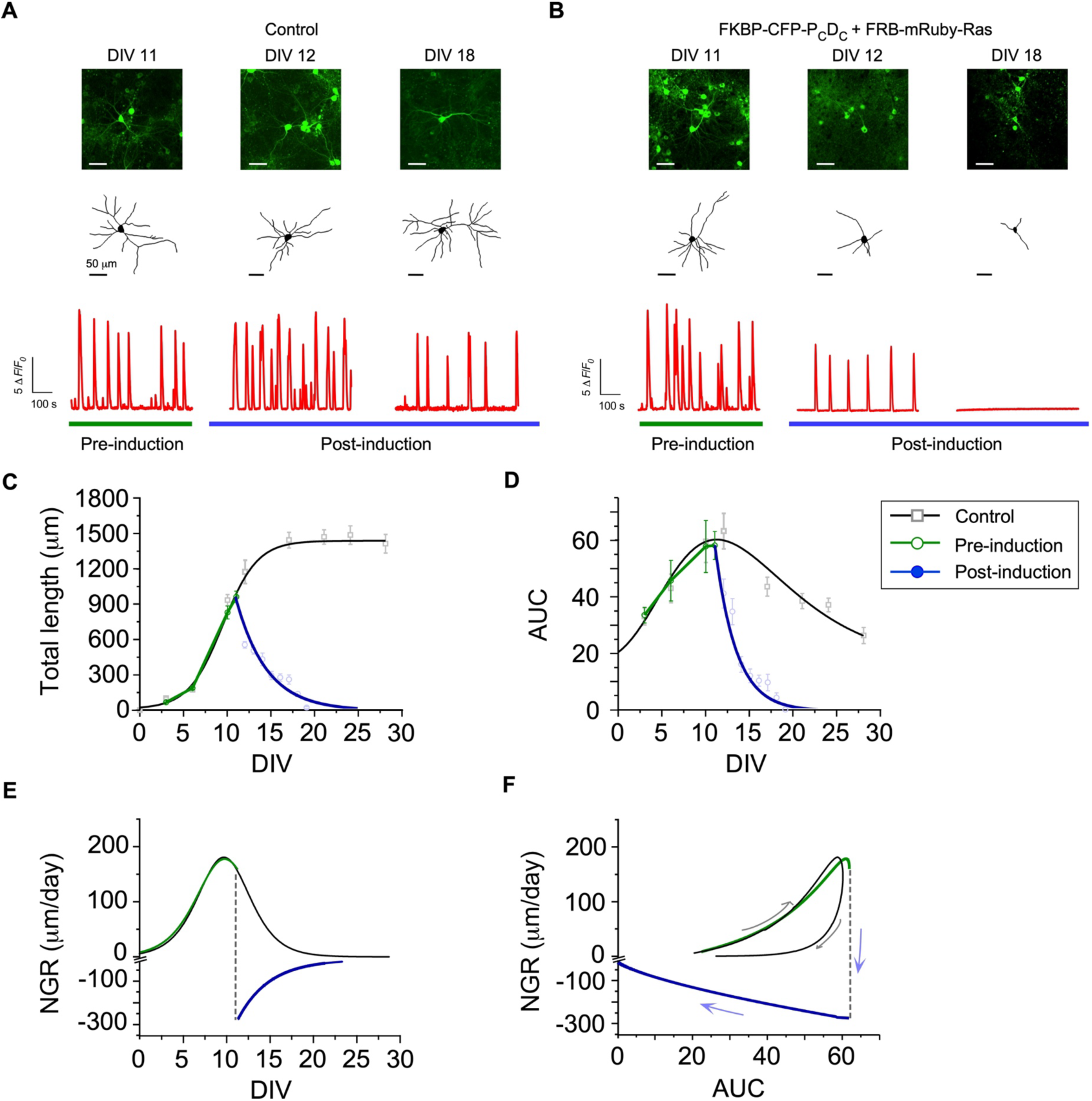
Inducible inhibition of the CaV1 influx-neuritogenesis coupling in cortical neurons. (A, B) Neurite tracing and fluorescence Ca^2+^ imaging, before (DIV 11) and after (represented by DIV 12 and DIV 18) 10 μM rapamycin induction, from the control neurons (A) versus the neurons expressing FRB-mRuby-Ras and FKBP-CFP-PCDC (B). (C) Time-dependent profiles of neurite outgrowth, described by the correlation between the total length per single neuron (μm) and the development stage (DIV), corresponding to the control neurons and the neurons subject to rapamycin-induced inhibition. (D) Time-dependent profiles of Ca^2+^ influx, described by the correlation between AUC (*ΔF*/*F0* per min) of Ca^2+^ oscillations and the development stage (DIV). (E) Temporal profiles of NGR (neurite growth rate, μm per day), which is derived from the total length-DIV curve (C). The dotted line represents the process of CMI induction. (F) Relationships between AUC (D) and NGR (E). The arrows indicate the temporal trends of neuronal development and its inhibition. All data points are calculated by mean ± SEM. See also Figure S10.

Our earlier data demonstrate that cultured cortical neurons go through an initial phase (∼14 days) of rapid outgrowth before entering into the plateau phase, forming a monotonic increasing curve (Figure 7C, sigmoidal-like curve in solid black), of which the first derivative indicates NGR of cultured cortical neurons (Figure 7E, bell-shaped curve in solid black). Inducing CMI by membrane association of PCDC led to reduction of Ca^2+^ oscillations, causing significant changes in neurite length, AUC and NGR, reflected as abrupt and large deviations from the curves of control neurons (Figure 7C-7E). Notably, the correlation between NGR of neuronal development and AUC of Ca^2+^ activities appeared to be a phase-plane plot that the trajectory for the control neurons was suddenly dragged down by CMI induction to the negative range of growth rate (Figure 7F).

Hippocampal neurons, well known for oscillatory activities, play vital roles in brain functions ^36^. We speculated that the CMI effects on the CaV1 activities-neuritogenesis coupling in cortical neurons should be generalizable, e.g., onto hippocampus. Membrane-anchored P-D peptides of Ras-YFP-PCDC significantly reduced oscillatory Ca^2+^ activity in hippocampal neurons, as quantified by AUC (Figure S9A and S9B). Meanwhile, the neurite length of hippocampal neurons expressing Ras-YFP-PCDC was significantly shorter than the control group of cytosolic peptides (Figure S9C and S9D).

## Discussion

In this study, we have developed a prototype of membrane-assisted CMI with opto-chemogenetic induction. It comprises NIR-responsive nanoparticles encapsulating rapamycin and peptides tagged with FRB/FKBP and Ras, encoding the DCT of CaV1.2 (P-D) or CaV1.3/1.4 (P+D) (Figure 8). Upon induction of membrane translocation, the peptides can form molecular-scale linkages to the targeted CaV1 channels, whether in long or short variants. Robust and potent inhibitory effects on CaV1 activities and neuronal development were acutely induced with ample dynamic range.

**Figure 8.**
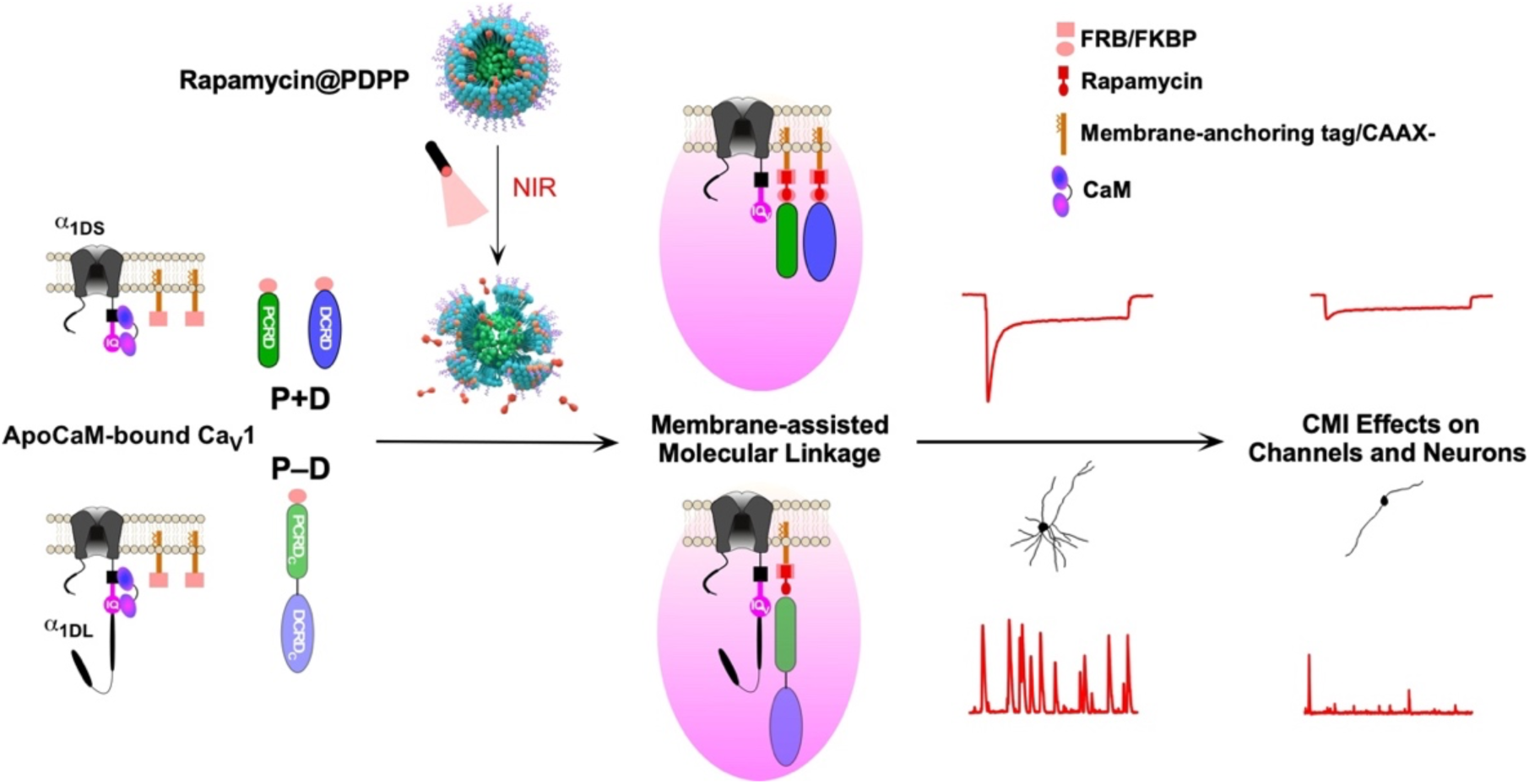
Summary of membrane-assisted molecular linkage and CaV1 inhibition with opto-chemogenetic induction. In neurons, CaV1 channels of two particular subgroups are targeted by CMI, represented by α1DS channels (without DCT) or a fraction of α1DL channels (containing the DCT domain but not auto-inhibited). This study designs and implements two major types of CMI peptides: P+D, represented by PCRD and DCRD peptides derived from CaV1.3/1.4 DCT; and P-D, represented by CaV1.2-encoded PCRD-DCRD. These peptides are devised mainly based on our data suggesting that the Ras/CAAX tags are able to form a type of effective linkage between the Ras-anchored peptides and the proteins on the membrane (indicated by pink shade). For α1DS, the P+D peptide has proven effective in creating a linkage; and for apoCaM-bound α1DL, P-D serves as the suitable solution, achieving minimum CMI at the basal state and potent CMI upon induction. The induction method is a combination of rapamycin-mediated FRB/FKBP binding and NIR-responsive rapamycin@PDPP nanoparticles, as our practical solution for opto-chemogenetics targeting CaV1. As proof of principle, P-D peptides are able to induce acute, specific and potent inhibitions on channels and neurons. This provides new evidence from a unique perspective, supporting the notion that CaV1-dependent Ca^2+^ oscillations and neurite outgrowth are tightly coupled.

### Insights into protein-protein interaction facilitated by membrane-anchoring

Besides ion channels, certain membrane proteins are known to interact with each other to form dimers or even oligomers, as the prerequisite of their subsequent signaling and activities ^25,37,38^. A prominent example is the Ras protein, whose dimerization on plasma membrane promotes RAF and MAPK cell signaling ^25^. In fact, the C-terminal domain containing CAAX as the membrane-anchoring tag in this study was adopted from H-Ras, one isoform of the Ras family ^14,31^. The G domain is critical to Ras dimerization, for which membrane-anchoring is also required in the first place ^39^. Our data, from both electrophysiology and FRET on the peptides tagged with Ras CAAX box, suggest that similar membrane-facilitated physical linkages and protein interactions may underlie Ras dimerization. This implies that CAAX-assisted translocation onto the membrane could increase the proximity of the molecules to a scale of 50 Å or less (Förster Radius), equivalent to a physical linkage (as required by our CMI principle) which enhances effective affinity or relative concentration (between intramolecular binding partners) and thus promotes the interaction/dimerization of the binding domains (e.g., G domain) ^40,41^. Another potential factor is via the lateral interactions between the transmembrane domains of membrane proteins ^42^, which, however, is unlikely the case of Ras/CAAX. Here, we quantitatively examined the equivalent peptide-channel dimerization using reliable biophysical methods (patch-clamp and 3-cube FRET) and mechanisms (DCT/CMI), which are readily expandable onto other membrane proteins including Ras. Interventions of Ras overactivation have been actively pursued as potential cancer therapeutics by targeting its membrane anchoring and dimerization, for which membrane-assisted CMI here may help develop (compound) screening assays, by taking advantage of the quantitative assays for CDI (electrophysiology) and Ca^2+^ dynamics (fluorescence imaging). The membrane-association tag of the tyrosine kinase Lyn and other c-Src family members is also important to dimerization and functions ^43–45^. We expect that Ras and Lyn (and potentially other membrane proteins) would share similar membrane (anchoring tag)-assisted linkage to enhance protein-protein interaction or dimerization, all awaiting future investigations to explore.

When we revised and finalized this manuscript, another study reported a similar strategy of utilizing the CAAX tag to demonstrate that Rad (small RGK G-protein) regulation of CaV1.2 channels is critically dependent on membrane-association ^46^. Ras tag was also proved by their FRET data to have the capability to bring the donor and acceptor close enough to yield high FRET efficiency. However, a few points are worth mentioning here. First of all, we reached our conclusion of the equivalent physical linkage based on the 3-component principle of CMI we established in earlier independent studies mainly with electrophysiology ^14,17^. Therefore, the core mechanism should be the membrane/Ras-assisted linkage in our view, leading to elevated effective concentration or enhanced apparent affinity as the consequences secondary to the formation of ‘physical’ linkage. In support, even Ras-tagged fluophores could produce high FRET in both our study (cellular FRET with membrane-anchored CFP/YFP) and their report (flowcytometric FRET with Cer/Ven). In addition, our experiments and analyses on membrane-assisted linkage, interaction and dimerization were restricted within the proteins or peptides on the membrane, different from their study heavily based on cytosolic/dispersed β2.

### Hints on mechanistic details of DCT, CaM and CMI

Before this work, only DCRD showed an appreciable (channel) affinity whereas the binding of PCRD/channel has not been detected yet, and thus the nature of their cooperativity still remains elusive. In the context of membrane-assisted linkage and CMI, membrane-anchored PCRD should also bind the channel when its effective concentration gets high enough, due to the fact that PCRD behaves similarly as DCRD (Figure 1C; Figure S5A; Figure S7A). Our data provide the missing piece of evidence critical to a unified mechanism of cooperative binding.

To date, the consensus holds that both the proximal and distal regions (PCRD and DCRD) are crucial for modulating CaV1 channels. For CaV1.2, PCRD and DCRD may have some electrostatic interactions according to *in silico* modeling ^30^, whereas for CaV1.3 no direct interaction could be detected between PCRD and DCRD ^14,19^. Later reports systematically comparing DCT across CaV1.1-1.4 demonstrate that a set of common principles are likely shared to induce inhibitory effects (CMI) on channel gating and Ca^2+^ influx ^14,17^. For instance, DCRD modules/motifs contribute more significantly than PCRD in CMI and related interactions. In support, FRET binding data suggest that DCRD itself could bind the CaM-binding IQ domain in contrast to PCRD that has no detectable binding; and PCRD and DCRD cooperate with each other to form the ternary complex of PCRD/DCRD/IQ ^14^. Meanwhile, the DCT peptide, as suggested, might behave as an apoCaM, the latter of which is known to bear intrinsic cooperativity between its two lobes ^47^. In light of this view and our data, one would postulate that PCRD by itself (just resembling DCRD) is also able to bind the channel/IQ, but with even lower affinity than DCRD. When present in the cytosol, the concentrations of PCRD or DCRD are way below the levels for any discernable binding with the channel to actually happen, thus no cooperative binding or CMI. However, once PCRD and/or DCRD translocate onto the membrane, the membrane-assisted linkage and the consequent (local) concentrations relative to each other (PCRD, DCRD and IQ/channel) are increased by orders of magnitudes to the levels of realistic binding. In this context, the observed cooperativity between PCRD and DCRD would resemble the two lobes of CaM: once one lobe/module is bound, the consequent allosteric changes facilitate the target binding of the other lobe/module, as the underlying principles for all the scenarios of P+D or P-D (e.g, Figure 1 and Figure 3). Not only to elucidate the molecular details of CMI/DCT but also to gain insights into the binding mechanisms related to CaM, the results and hints from this study merit further investigation.

### The development of molecular tools for protein dimerization

Controllable protein heterodimerization allows to modulate diverse biological processes such as protease activity, transcription and translocation ^48–51^, thus applicable to a broad spectrum of scenarios as powerful molecular tools ^52,53^. Rapamycin-inducible FKBP/FRB heterodimerization has been widely applied due to its ultra-high affinity of rapamycin towards its protein binding partners FKBP and FRB (*Kd* = 12 nM for FKBP-rapamycin-FRB), with a clear baseline (no detectable binding) in the absence of rapamycin ^54,55^. To improve the spatial control of rapamycin system, photocaged rapamycin analogues have been developed, such as cRb, which features a nitrobenzyl cage linked to a rapamycin analogue ^56^, dRap, a photo-cleavable rapamycin dimer ^57^, pRap, a caged rapamycin with nitro-piperonyloxycarbonyl N-hydroxysuccinimide carbonate (NPOC-NHS) ^58^, DMNB caged rapamycin ^59^, and arylazopyrazole rapamycin analogs ^60^. However, all of these photocaged rapamycin variants are triggered by the ultraviolet light (UV), unfavored by biological applications. In fact, most approaches for optical control of protein-protein interaction by photocaged drugs are limited by the short wavelength stimulation ^61,62^. In comparison, NIR light has less cell damage and deeper tissue penetration. Moreover, unlike rapamycin analogues, our design of rapamycin@PDPP does not change the skeleton of rapamycin, thus relieving the concerns regarding potential reduction in membrane permeability and increase in cellular toxicity due to covalent modifications of rapamycin. And nanoparticles are recognized as a well-established platform for controlled drug delivery, which promises extra benefits for *in vivo* applications ^32^.

Besides opto-chemogenetics based on rapamycin@PDPP and FKBP/FRB in this work, other optically-controlled dimerization methods (such as CRY2/CIB, UVR8/UVR8, and light-switchable nanobodies) are commonly triggered by blue light or UV light ^63–65^. Some newly designed optogenetic systems could be triggered by NIR light but suffer from low efficacy when compared to the FKBP/rapamycin/FRB system in practice ^66,67^. Here in this work, by taking advantage of two separate but established lines of work, we designed and implemented a prototype of NIR-triggered heterodimerization, as a feasible strategy of optogenetics.

Moreover, the membrane-assisted molecular linkage as proposed in this work opens up new avenue to design dimerization tools. The first step is to adopt or devise cytosolic modules with no or low weak affinities to dimerize; then, the cytosolic modules need to translocate onto the membrane, e.g., by attaching membrane-anchoring tags such as Ras CAAX to monomeric proteins or peptides. Combining with other molecular tools (such as rapamycin@PDPP in this work), a general platform with controllable binding/dimerization can be achieved. It is a challenging task to specifically design and implement optical control within each particular molecule, considering the diverse interactions and mechanisms often difficult to intervene/control. Instead, through a more universal strategy as demonstrated in this work, the desired dimer or even binding complex can be achieved on the membrane with high spatiotemporal resolutions without having to dig into the binding details.

### Insights into the coupling of CaV1 activities with neuronal development

Cellular Ca^2+^ signals, especially transmembrane Ca^2+^ influx, play a central role in neural development ^35,68^. Spontaneous, regenerative and correlated or even synchronized Ca^2+^ activities are particularly important to neurite outgrowth, presumably owing to enhanced efficiency in gene transcription in the nucleus and related downstream events ^69^. In addition, L-type Ca^2+^ channels are proposed as the core mediator to specifically and mechanistically link Ca^2+^ oscillations, the CaMKII-CREB signaling pathway, and also neuritogenesis altogether, i.e., the CaV1 activity-neuritogenesis coupling ^17^. However, due to multiple complications, this coupling needs additional evidence before it can be fully established. The first complication is that transmembrane Ca^2+^ has an array of sources, e.g., N-Methyl-D-aspartate (NMDA) receptors, TRP channels, Orai/STIM, and CaV2 ^70^. Another complication comes from the molecular tools that most studies are relying on: pharmaceutical interventions based on DHP derivatives. Both CaV1 inhibitors and potentiators have generated the data supporting the proposed coupling, but also encountered with many complicated scenarios, e.g., the unexpected neuroprotection by DHP inhibitors, the differential effects of Cd^2+^ and nimodipine on transcription signaling, and the off-target effects and neurotoxicity of Bay-K-8644 ^71–73^. Moreover, knock-out mice of CaV1.3-/- and CaV1.2-/- could also provide useful information ^69,74^; however, they may not be suitable to certain studies on neural development due to compensatory effects as reported ^75,76^. CMI peptides stand out as a promising toolset to explore the molecular physiology of CaV1 ^17^. Now in this study, with optimized P-D peptides and membrane-assisted inducible CMI, unprecedented evidence has been provided to quantitatively characterize the coupling and its modulation. In this work, we not only validate our design in both recombinant and neuronal systems, but also provide the direct evidence strongly supporting that (apoCaM-bound) CaV1 is the molecular basis of the oscillatory Ca^2+^ signals in tight coupling with neurite outgrowth in cultured cortical neurons (Figure 7). We expect that future studies deploying the findings and tools from this study will greatly help elucidate the signaling pathways, mechanistic details and relevant pathophysiology of CaV1 in neurons.

### Therapeutic potentials of opto-chemogenetics and rapamycin@PDPP nanoparticles

The proof-of-concept of opto-chemogenetic CMI highlights the therapeutic potential of CMI-based antagonism in CaV1-related disorders, such as Parkinson’s disease ^14^. In fact, some subtypes of congenital long QT syndromes are genetically associated with CaV1 and CaM, both of which lead to alternations in channel activities and severe cardiac arrhythmias ^77,78^. Thus, CaV1 antagonisms based on CMI may serve as the candidate treatments for LQT or like diseases. Moreover, NIR-controlled CMI may enable noninvasive, high spatiotemporal, and tissue compatible interventions of CaV1 in the heart and the brain. Notably, our design targets CaV1.3 as the representative subtype, presumably expandable on CaV1.2 due to their shared CMI effects and mechanisms ^17^. Multiple variants and isoforms are expressed in the brain and heart ^20,22,23,79–82^. As demonstrated, P+D and P-D peptides have differential capabilities to deal with CaV1.3 variants, which include both the long and short α1D in cortical and hippocampal neurons (Figure 8). For any particular type of primary cells, it is necessary to conduct tests and analyses on each channel variant with different peptides before proceeding further. CaV1.2-encoded P-D peptides derived from this work have displayed excellent performance and characteristics, which invite additional optimizations and further explorations with healthy and diseased neurons.

Enlightened and encouraged by optogenetics (e.g., channelrhodopsin), an increasing number of optical tools have been developed for precise spatiotemporal control of physiological processes in living cells, which are designated as ‘optophysiology’ ^83^. Optically-controlled and genetically-encoded Ca^2+^ channel actuators/modulators are promising research directions in ‘optophysiology’ ^84^. In parallel, chemogenetics provides potent and robust control of physiological processes ^85^. Development of chemogenetic Ca^2+^ channel modulators has also been actively pursued to facilitate both basic and therapeutic research ^31^. Our NIR-triggered CMI has seamlessly integrated rapamycin@PDPP with chemogenetics, offering at least three aspects of advantages: 1. effective and robust control of FRB/FKBP binding (rapamycin); 2. high precision of spatiotemporal control at single-cell resolution (photosensitive nanoparticles); 3. deep tissue penetration with long-wavelength photostimulation (NIR) and high biocompatibility (nanomaterials and genetically-encoded modulators). With all these benefits including the unique feature of NIR ^86,87^, we expect that our opto-chemogenetics prototype would be applicable broadly and particularly beneficial to future *in vivo* applications.

## Author Contributions

X.D.L. conceived, designed and supervised the project; Y.B.F. and C.F.X. helped organize and oversee the project. J.L.G., Y.X.Y., B.Y.L., Z.Y., S.Q., W.Z., S.X.G., Y.L. and B.W. performed the experiments and analyzed the data. N.L. provided preliminary patch-clamp data on peptide effects. X.D.L. wrote and finalized the manuscript. All authors contributed to writing and revising the manuscript

## Acknowledgments

We thank all X-Lab members for discussions and help. This work is supported by the grants from the National Natural Science Foundation of China (81971728 and 22077025), the Natural Science Foundation of Hebei Province (B2023202030) and the China Postdoctoral Science Foundation (2023M740968).

## Declaration of interest

The authors declare no competing interests.

## Methods

### Molecular biology

YFP-FKBP-DCRD (Addgene ID: 87453), YFP-FKBP-PCRD (Addgene ID: 87452), and FRB-CFP-Ras (Addgene ID: 87451) are available on Addgene. YFP-PCRD, YFP-DCRD, CFP-DCRD, YFP-CCATC, CaV1.3 α1DS (AF370009.1) and CaV1.3 α1DL (NM_001389225.2) are from previous study^14,17^. PCRD-CFP-Ras and DCRD-CFP-Ras were generated by replacing the FRB in FRB-CFP-Ras with PCRD or DCRD, respectively. Constructs of PCRD-YFP-Ras, CFP-Ras, YFP-Ras and CFP-YFP-Ras were made by appropriate design. For CFP-ER/K-YFP-Ras, ER/K α-helix (206 a.a.) were inserted between CFP and YFP sequence of CFP-YFP-Ras with BspEI sites ^29^. For Ras-mRuby-CCATC, the Ras tag (KLNPPDESGPGCMSCKCVLS) was PCR-amplified and fused to the N-terminus of mRuby, then inserted into YFP-CCATC with KpnI and NotI to replace YFP. For FKBP-YFP-PCDC, the FKBP was PCR-amplified with KpnI and BamHI sites and fused to the N-terminus of YFP-PCDC and then cloned into pcDNA4 vector. Ras-FKBP-YFP-PCDC, with PCR-amplified Ras tag was fused to the N-terminus of FKBP-YFP-PCDC, inserting into a customized pcDNA3 vector via unique KpnI and XbaI sites.

### Transfection of cDNA constructs in HEK293 cells

HEK293 cell line (ATCC) was free of mycoplasma contamination, checked by PCR with primers 5’-GGCGAATGGGTGAGTAACACG - 3’ and 5’- CGGATAACGCTTGCGACCTATG -3’. For electrophysiology recording, cells were cultured in 60 mm dishes. Recombinant channels were transiently transfected according to an established calcium phosphate protocol ^19^. We applied 4 μg-5 μg of cDNA encoding the desired channel α1D subunit, along with 4 μg of rat brain β2a (NM_053851.2) and 4 μg of rat brain α2δ (NM_012919.3) subunits. Additional 2 μg of cDNA was added as required in co-transfections. cDNA for simian virus 40 T antigen (1 μg) was also co-transfected to enhance the expression of channels. Cells were washed with PBS 6 hours after transfection and maintained in culture medium of supplemented DMEM, then incubated for at least 48 hours in a water-saturated 5% CO2 incubator at 37°C before whole-cell recordings. For confocal fluorescence imaging, HEK293 cells were cultured in 35 mm confocal dishes. 2 μg of desired cDNA was transfected by lipofectamine 2000 (Invitrogen) for 6 hours. Cells were used after 2 days.

### Whole-cell electrophysiological recording

Whole-cell recordings of transfected HEK293 cells were obtained at room temperature (25°) using an Axopatch 700B amplifier (Axon Instruments). Electrodes were pulled with borosilicate glass capillaries by a programmable puller (P-1000, Sutter Instruments, Novato, CA) and heat-polished by a microforge (MF-830, Narishige, Japan), resulting in 3-5 MΩ resistances, before series resistance compensation of about 70%. The internal solutions contained (in mM): CsMeSO3, 135; CsCl, 5; MgCl2, 1; MgATP, 4; HEPES, 5; and EGTA, 5, adjusted to 290∼300 mOsm with glucose and pH 7.3 with CsOH. The extracellular solutions contained (in mM): TEA-MeSO3, 135; HEPES, 10; CaCl2 or BaCl2, 10, adjusted to 300∼310 mOsm with glucose and pH 7.3 with TEAOH. Whole-cell currents were generated from a family of step depolarization (-60 to +50 mV from a holding potential of -70 mV) or a series of repeated step depolarization (-10 mV from a holding potential of -70 mV). Currents were recorded at 2 kHz low-pass filtering. Traces were acquired at a minimum repetition interval of 30 s. P/8 leak subtraction was used throughout. Rapamycin (Solarbio, or Aladdin) was dissolved in DMSO as 10 mM or 1 mM stock solution, stored at -20°C, and then diluted to 1 μM using extracellular Ca^2+^ solution before electrophysiological recordings.

### 2-hybrid 3-cube FRET

All the FRET experiments were performed in Tyrode’s buffer containing 2 mM Ca^2+^. An inverted epi-fluorescence microscope (Ti-U, Nikon) was used with computer-controlled filter wheels (Sutter Instrument) to coordinate with dichroic mirrors for appropriate imaging at excitation, emission, and FRET channels. The following filters sets were utilized: excitation: 438/24 nm and 480/30 nm; emission: 483/32 nm and 535/40 nm; dichroic mirrors: 458 nm and 505 nm. Fluorescence images were acquired with a Neo sCMOS camera (Andor Technology), which were analyzed with 3^3^-FRET algorithms coded in Matlab (Mathworks).

### Confocal fluorescence imaging

Fluorescence images were obtained in HEK293 cells expressing membrane-localized CFP-tagged FRB and with YFP-tagged cytoplasmic FKBP-PCRD/DCRD on 30 mm confocal dishes. The images were captured at 30 s intervals. Cells were applied with 1 μM rapamycin, 0.1% DMSO (vehicle), and 25 μg/ml rapamycin@PDPP with or without the stimuli of 808 nm near-infrared laser using a Hi-Tech high power laser generator. Images were recorded with Olympus Fluoview FV300 or Zeiss LSM710 laser scanning confocal microscopes. Images were analyzed using Imaris 7.7.2 and ImageJ. Fluorescence imaging of cultured neurons was performed on Dragonfly High Speed Confocal Microscope (Dragonfly 200, Andor, England) and with Fusion software. Measurement of the total length for neurites was performed with Imaris 7.7.2 (Bitplane). Only non-overlapping neurons were selected for analysis and images of at least 21 neurons from two independent culture preparations were analyzed.

### Preparation of drugs@PDPP nanoparticles

PDPP (0.5 μμ), DSPE-PEG2000-NH2 (3 mg, Tansh-Tech Technology Company, Guangzhou, China) and rapamycin (0.5 mg) were dissolved in 1 ml of THF (tetrahydrofuran) and sonicated for 30 min, then DPPC (8 mg, Rowen) was dissolved in 200 μl of dichloromethane and sonicated. The two solutions were mixed and sonicated again for 30 min, and the mixture was quickly transferred to 9 ml of ultrapure water until the ultrasonic solution became clear, and then placed on a stirring table to stir for 8 hours. Argon was blown into the solution for 1.5 hours, and then the solution was placed on a stirring table to stir for 8 hours to obtain the rapamycin@PDPP solutions. The preparation of Cy5@PDPP solutions was similar to rapamycin@PDPP solutions. Briefly, rapamycin was not added to THF at the first step. The mixture solution containing PDPP, DSPE-PEG2000-NH2 and DPPC was heated to 65° and then added with Cy5. After stirring for 10 min, the solution was ultrasonic for 30 min to get Cy5@PDPP solution. In addition, the solution was transferred to a 3500k dialysis bag, and the excess organic solvent was removed by dialysis for two days to obtain the final rapamycin@PDPP or Cy5@PDPP nanoparticles. For Cy5&rapamycin@PDPP nanoparticles, rapamycin@PDPP solution was mixed with Cy5 (0.5 mg), heated to 65°, stirred for 10 min, and sonicated for 30 min to obtain the Cy5&rapamycin@PDPP nanoparticles. All nanoparticle solutions were centrifuged at 6,500 rpm for 20 min to obtain concentrated solutions after ultrafiltration, and the concentration of nanoparticles was determined by ultraviolet absorption spectrum for subsequent experiments.

### Determination of UV absorption spectra of drug@PDPP nanoparticles

Rapamycin@PDPP, Cy5@PDPP, and Cy5&rapamycin@PDPP nanoparticles were concentrated to a concentration greater than 500 μg/ml using a 100k ultrafiltration tube. The concentrated solution of 30 μl nanoparticles was diluted 20 times, then the sample was scanned by UV absorption spectrum. The UV absorbance of nanoparticles at 808 nm was recorded to calculate the concentration according to the standard curve equation of PDPP.

### Particle size measurement of drug@PDPP nanoparticles

Ultrapure water was used to dilute the nanoparticles to an appropriate concentration for reserve. The dynamic light scattering particle size meter was turned on and preheated for 20 min in advance. 1 ml of the nanoparticle solution was added to the dedicated particle size colorimeter and put into the detection tank to measure the particle sizes.

### Analysis of temperature rise curve of rapamycin@PDPP nanoparticles

200 μl of ultra-pure water was added to a 96-well plate, and the temperature of the solution was recorded under near-infrared laser irradiation (808 nm, 5 min, Hi-Tech high power laser generator). Subsequently, rapamycin@PDPP nanoparticles with concentrations of 5 μg/ml, 10 μg/ml and 20 μg/ml were irradiated according to the requirements of sample addition and irradiation, and the change in solution temperature was recorded. The obtained data were imported into the drawing software for analysis, and the temperature rise statistics of nanoparticles were completed.

### Photothermal stability analysis of rapamycin@PDPP nanoparticles

Rapamycin@PDPP (200 μl) nanoparticles were added to the 96-well plate and irradiated with a laser at 808 nm for 5 min to achieve the highest temperature. Then the laser was turned off to allow cooling for 15 min to return to its original temperature. Real-time readings were taken every 30 seconds during heating and cooling cycles. This procedure was repeat 3 times to complete the whole cycle of heating and cooling.

### Near-infrared photothermal imaging analysis of rapamycin@PDPP nanoparticles

Rapamycin@PDPP (200 μl) nanoparticles were placed in a 96-well plate, and the nanoparticles solution was irradiated by a laser at 808 nm. Photos were taken by a near-infrared imager (FLIR T420 infrared thermal imager camera) every 60 s, and the images were processed by data processing software to complete the near-infrared thermal imaging analysis of nanoparticles.

### Confocal fluorescence imaging of Cy5 released from Cy5&rapamycin@PDPP nanoparticles

2 ml of prepared nanoparticles were dropped onto confocal dishes and irradiated with 808 nm laser for 0, 0.5, 1, 2, 3, 4 or 5 min, respectively. Then samples were dried and imaged with a Leica SP5 confocal laser scanning microscope to capture the fluorescence of Cy5.

### Dissection and culturing of cortical and hippocampus neurons

Cortical neurons or hippocampus neurons were dissected from newborn ICR mice or SD rat, respectively. The tissues of cortex or hippocampus were isolated and then digested with 0.25% trypsin without EGTA for 15 min at 37°C. Then digestion was terminated by Dulbecco’s modified Eagle medium (DMEM) supplemented with 10% fetal bovine serum (FBS) and 1% antibiotics. The cell suspension was sieved through a filter and centrifuged at 1,000 rpm for 5 min. The cell pellet was resuspended in DMEM supplemented with 10% FBS and were plated on poly-D-lysine-coated 35 mm No. 0 confocal dishes (In Vitro Scientific). After 4 hours, neurons were maintained in Neurobasal medium supplemented with 2% B27 and 1% GlutaMAX-I (growth medium), and cultured in the incubator with temperature of 37°C and 5% CO2. All animals were obtained from Beijing Vital River Laboratory Animal Technology Co., Ltd. Procedures involving animals were approved by local institutional ethical committees (Beihang University).

### Virus infection on cultured neurons

AAV2/DJ-*hSyn*-jGCaMP7b-XC virus was used for infection of cultured neurons (Hanbio Biotechnology, China). Other viruses include pSLenti-*hSyn*-FKBP-CFP-PCDC-WPRE, pSLenti-*hSyn*-FRB-mRuby-Ras-WPRE and pAAV2/9 -*hSyn*-NES-jRGECO1a-WPRE viruses (OBiO Technology, China). 1 µl of 1 × 10^12^ v.g./ml of the desired adeno-associated virus or 1-2 µl of 1 × 10^8^ TU/ml of the desired lentivirus were added to growth medium at DIV 0 unless otherwise indicated. Neuronal experiments were repeated independently at least twice.

### Transfection of cDNA constructs in neurons

1 μg of cDNA mixed with 1μl PLUS^TM^ reagent were transiently transfected into DIV 3-7 cultured neurons by Lipofectamine™ LTX (Invitrogen) with a typical protocol according to the manual. The Opti-MEM containing plasmids and Lipofectamine™ LTX was added to the Neurobasal medium for transfection. After 2 hours, neurons were maintained in Neurobasal medium supplemented with 2% B27 and 1% GlutaMAX-I for at least 2 days before imaging.

### Data analysis and statistics

Data were analyzed in Matlab, OriginPro and GraphPad Prism software. The values of standard error of mean (S.E.M) or standard derivation (S.D.) were calculated. Two-tailed Student’s *t*-test, paired *t-*test, or One-way ANOVA followed by Dunnett for post hoc test were applied when applicable. *, *p*<0.05; **, *p*<0.01; ***, *p*<0.001; n.s. or not significant, *p*>0.05.

## Notes

### Competing Interest Statement

The authors have declared no competing interest.

