## Supplementary Figures for "Opto-chemogenetic inhibition of L-type Ca_V_1 channels in neurons through a membrane-assisted molecular linkage"

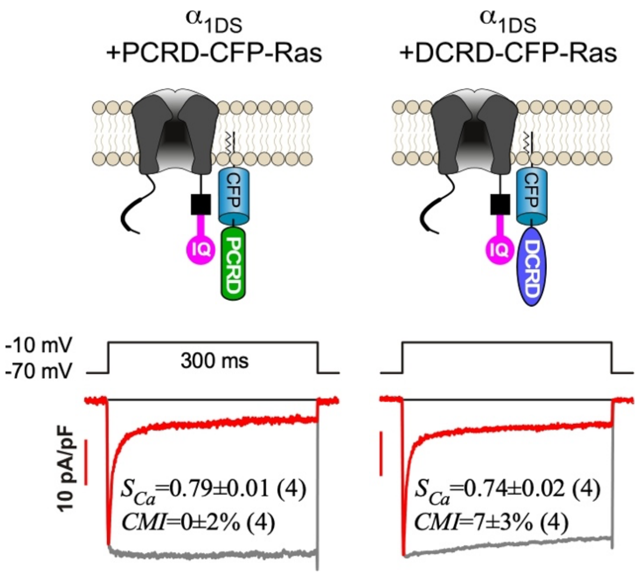


**Figure S1. Membrane-anchored PCRD or DCRD alone failed to inhibit $\alpha_{1DS}$.**

The cartoon illustration of PCRD (left) or DCRD (right) on the membrane anchored by Ras/CAAX. Representative current traces ($I_{Ca}$, red and $I_{Ba}$, gray) are shown. The levels of CDI and CMI are quantified by $S_{Ca}$ and *CMI*, respectively.

All statistical data are given as mean ± SEM.

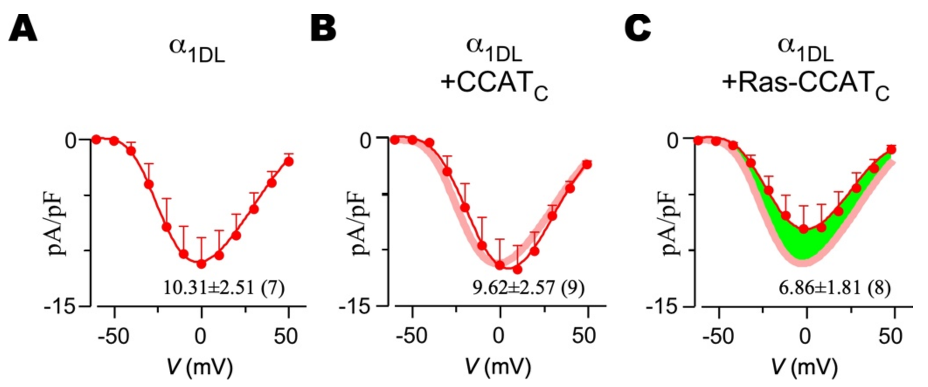


**Figure S2. Profiles of the effects on $Ca^{2+}$ current amplitudes of $\alpha_{1DL}$ by cytosolic or membrane-anchored P-D peptides.**

(A-C) The profiles of $Ca^{2+}$ current density (pA/pF) are in accordance with Figure 3A-3C. The profile of the $\alpha_{1DL}$ control (A) was replotted (in pale red) to compare with the cytosolic P-D (YFP-$CCAT_C$, B), and membrane-anchored P-D (Ras-mRuby-$CCAT_C$, C). The values of the peak current density ($I_{peak}$, pA/pF) at -10 mV are labelled for each group. The green shade (C) illustrates the difference in the voltage-dependent profiles between the Ras-tagged and untagged groups.

All statistical data are given as mean ± SEM.

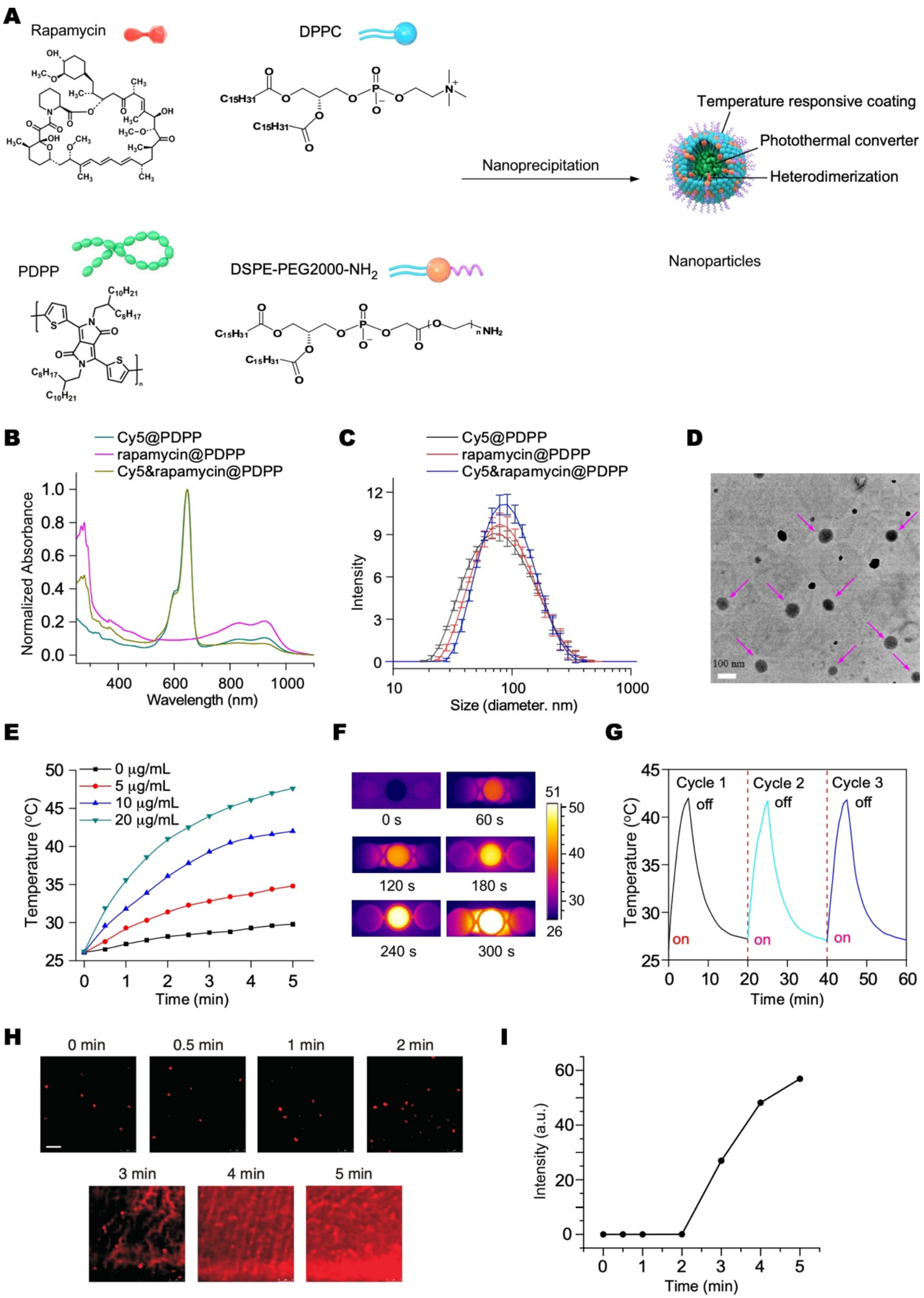


**Figure S3. Fabrication and characterization of the nanoparticles with optical control of rapamycin release.**

(A) Schematic illustration of the process to prepare PDPP nanoparticles encapsulating rapamycin (rapamycin@PDPP). Rapamycin@PDPP was prepared by nano-precipitation. PDPP plays a photothermal role, DPPC and DSPE-PEG2000-$NH_2$ are lipid coatings, and rapamycin is a drug molecule.

(B) Normalized absorption spectra of Cy5@PDPP, rapamycin@PDPP, and Cy5&rapamycin@PDPP nanoparticles, measured by the UV absorption spectrum.

(C) Sizes (in diameter) of Cy5@PDPP, rapamycin@PDPP, and Cy5&rapamycin@PDPP nanoparticles dispersed in water, measured by dynamic light scattering particle size distribution.

(D) TEM image of rapamycin@PDPP. Red arrows indicate intact nanoparticles. The scale bar is 100 nm.

(E) Temporal profiles of the temperature of solutions containing different concentrations of rapamycin@PDPP under near-infrared (NIR) laser irradiation (808 nm, 1 W $cm^{-2}$, 5 min). The concentration of rapamycin@PDPP nanoparticles is calibrated according to the ultraviolet standard curve of conjugated polymer PDPP.

(F) Infrared thermal images of the aqueous solution containing 25 $\mu$g/ml rapamycin@PDPP collected at different laser irradiation times as marked below. Color bar represents the temperature. Only the middle well contains the solution of rapamycin@PDPP.

(G) Time-temperature profiles of three cycles of photothermal heating and natural cooling to test the stability of rapamycin@PDPP nanoparticles. The temperature was monitored in real time during the experiments.

(H) Confocal fluorescent images of Cy5 (colored in red) releasing from the Cy5&rapamycin@PDPP nanoparticles upon different laser irradiation times. Solutions containing nanoparticles were dropped on the confocal dishes, and naturally dried after laser irradiation for confocal imaging.  The scale bar is 500 nm.

(I) The temporal profile of Cy5 releasing from Cy5&rapamycin@PDPP nanoparticles upon laser irradiation. The fluorescence of background without the light spots of nanoparticles was calculated and summarized.

All statistical data are given as mean ± SD (Standard Deviation).

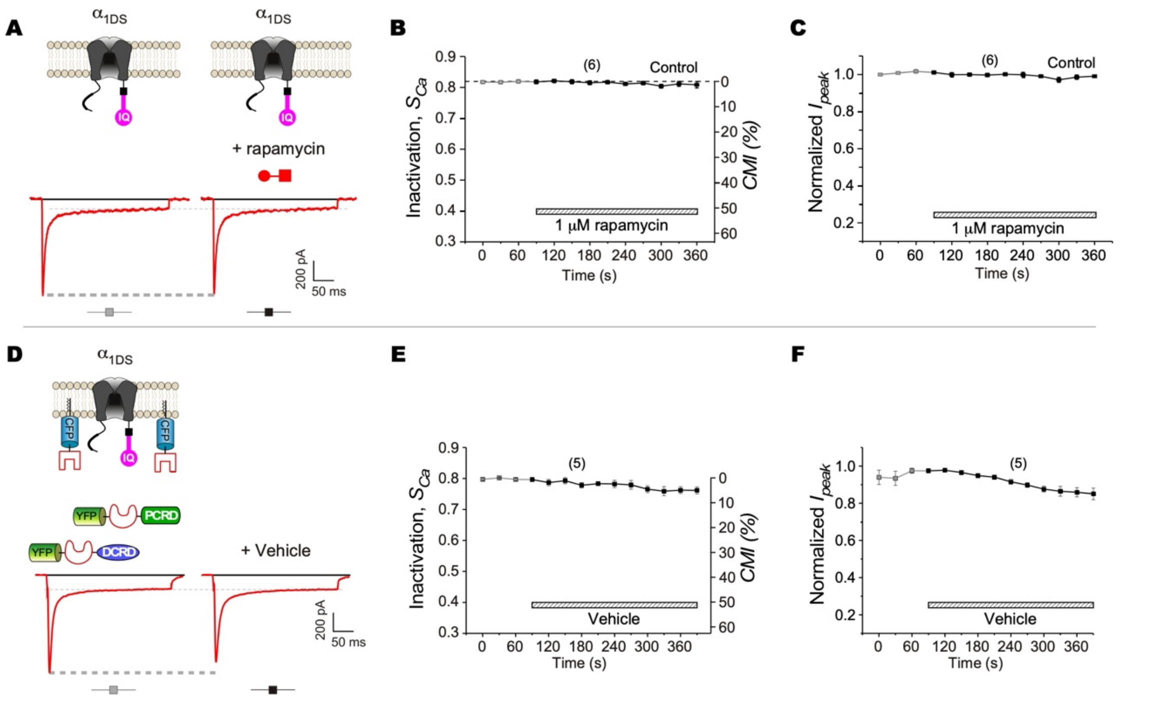


**Figure S4. Negative control of chemo-genetically inducible CMI on $\alpha_{1DS}$.**

(A-C) Rapamycin caused no apparent effect on control group of $\alpha_{1DS}$ channels (without P+D expression). Exemplars of $Ca^{2+}$ current traces demonstrate no changes in $Ca^{2+}$ currents by 1 μM rapamycin, where both the peak ($I_{peak}$) and steady-state amplitude (at 300 ms, i.e., $I_{300}$) are indicted by the dashed lines (A). Temporal profiles of $S_{Ca}$ along with *CMI* (B) and normalized $I_{peak}$ (C) are shown. Data points connected by gray and black lines represent the values of $S_{Ca}$ or $I_{peak}$ before and after rapamycin application, respectively.

(D-F) $Ca^{2+}$ currents were not affected by the vehicle (0.1% DMSO). HEK293 cells expressing recombinant $\alpha_{1DS}$, FRB-CFP-Ras, YFP-FKBP-PCRD and YFP-FKBP-DCRD (P+D) are treated with vehicle (0.1% DMSO) as the negative control. Exemplars of $Ca^{2+}$ current traces demonstrate no affected by 0.1% DMSO featured with a characteristic stable of $I_{peak}$ and $I_{300}$ (D). Temporal profiles of $S_{Ca}$ along with potency *CMI* (E) and normalized $I_{peak}$ (F).

All statistical data are given as mean ± SEM.

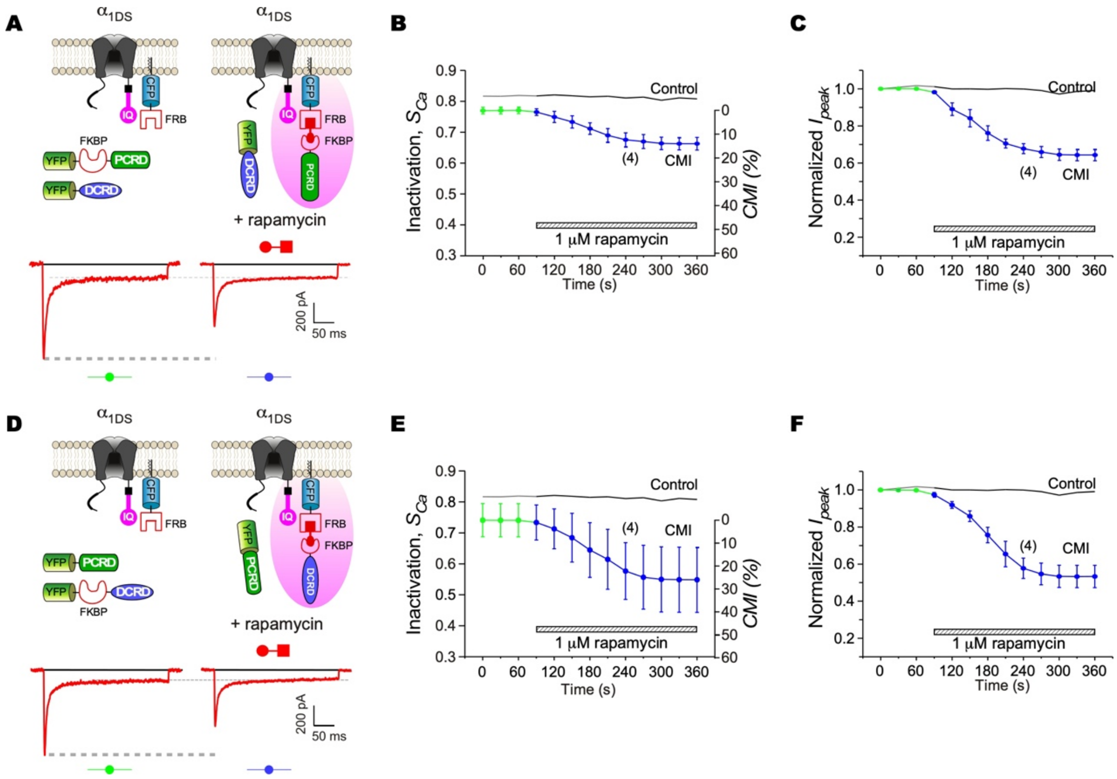


**Figure S5. Rapamycin-induced membrane-assisted CMI on $\alpha_{1DS}$.**

(A-C) Rapamycin-induced CMI through translocation of PCRD onto the membrane. Membrane-anchored FRB (FRB-CFP-Ras) recruited YFP-FKBP-PCRD to the membrane by rapamycin-mediated FRB/FKBP binding. Exemplars of $Ca^{2+}$ current traces demonstrate the effective inhibition by 1 μM rapamycin, where both the peak ($I_{peak}$) and steady-state amplitude ($I_{300}$) are indicted by the dashed lines (A). Temporal profiles of $S_{Ca}$ along with *CMI* (B) and normalized $I_{peak}$ (C) demonstrate rapamycin-induced attenuation in comparison with $\alpha_{1DS}$ control. Data points connected by lines (in green and blue) represent the values of $S_{Ca}$ or $I_{peak}$ before and after rapamycin application, respectively.

(D-F) Rapamycin-induced CMI through translocation of DCRD onto the membrane. Similar to the first scheme, YFP-FKBP-DCRD is mobilized to the membrane by rapamycin to form the effective connection between $\alpha_{1DS}$ and DCRD. Exemplars of $Ca^{2+}$ current traces demonstrate effective inhibition by 1 μM rapamycin, where both $I_{peak}$ and $I_{300}$ are indicted by the dashed lines (D). Temporal profiles of inactivation $S_{Ca}$ along with potency *CMI* (E) and normalized $I_{peak}$ (F). All statistical data are given as mean ± SEM.

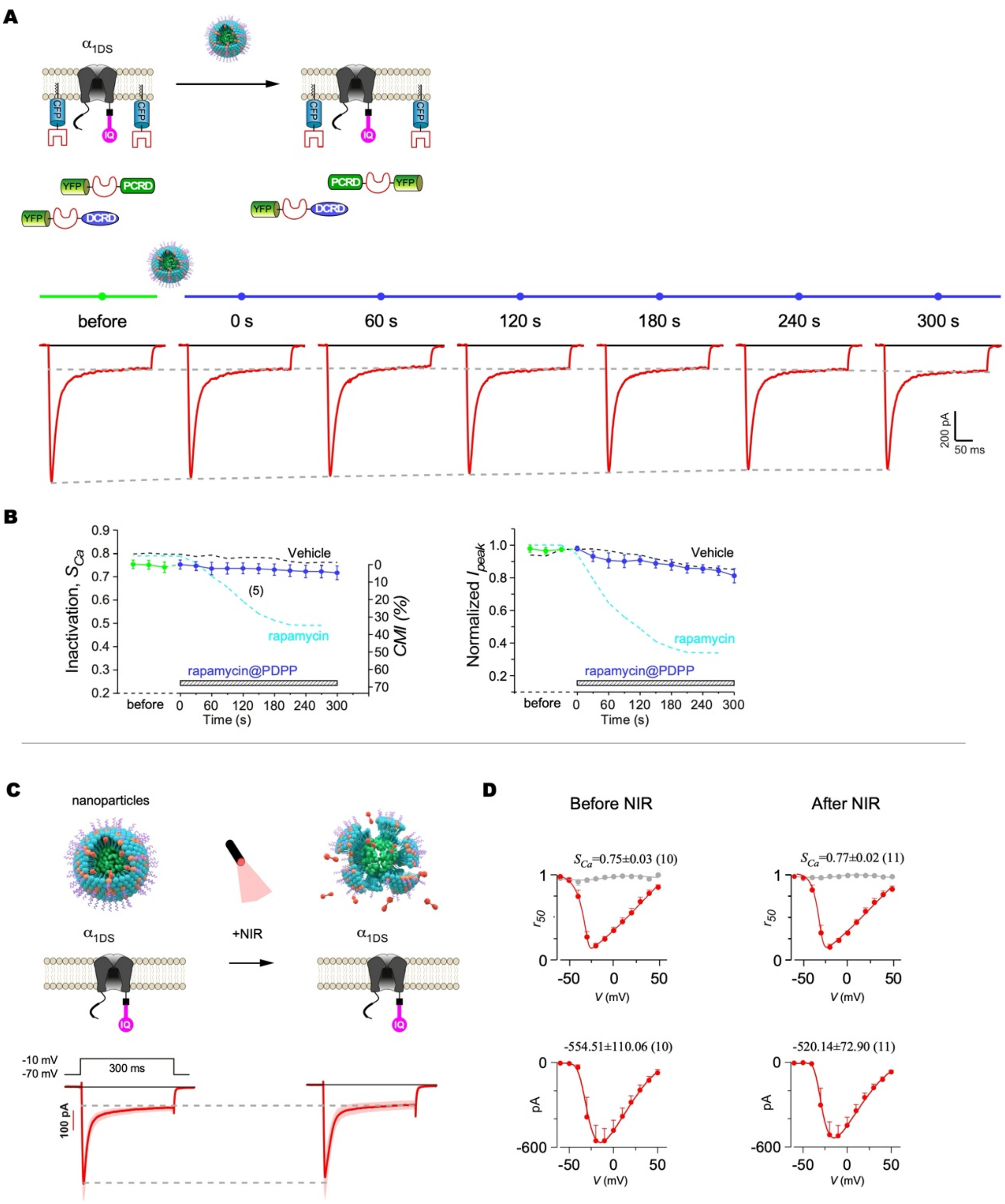


**Figure S6. Negative controls for opto-chemogenetic CMI on $\alpha_{1DS}$.**

(A, B) Negative control of opto-chemogenetic CMI for $\alpha_{1DS}$. HEK293 cells expressing recombinant $\alpha_{1DS}$, FRB-CFP-Ras, YFP-FKBP-DCRD and YFP-FKBP-DCRD were treated with rapamycin@PDPP nanoparticles without NIR stimuli. Representative $Ca^{2+}$ traces at different timepoints exhibited stable $I_{peak}$ and $I_{300}$ (A). Temporal profiles of $S_{Ca}$ or *CMI* (B, left) and normalized $I_{peak}$ (B, right). Black dotted curves represent the treatment with vehicle (0.1% DMSO) as the negative control (adapted from Figure S4D-S4F), while cyan dotted curves correspond to rapamycin treatment as the positive control (adapted from Figure 5C).

(C, D) Profiles of $\alpha_{1DS}$ currents effected by rapamycin@PDPP. Cells expressing $\alpha_{1DS}$ (without P+D expression) were treated with rapamycin@PDPP nanoparticles. NIR irradiation for 5 min to trigger the release of rapamycin reaching the final concentration of 1 μM, then recorded with full range of *V*. $I_{Ca}$ trace (averaged) of $\alpha_{1DS}$ at the membrane potential (*V*) of -10 mV before or at the ending points of NIR stimuli *in situ* (C). Voltage-dependent profiles of $r_{50}$ and $I_{peak}$ (pA) of the channels are shown before NIR or treated NIR at the ending timepoints (D). The values of $S_{Ca}$ and $I_{peak}$ at -10 mV are indicated alongside the plots.

All statistical data are given as mean ± SEM.s

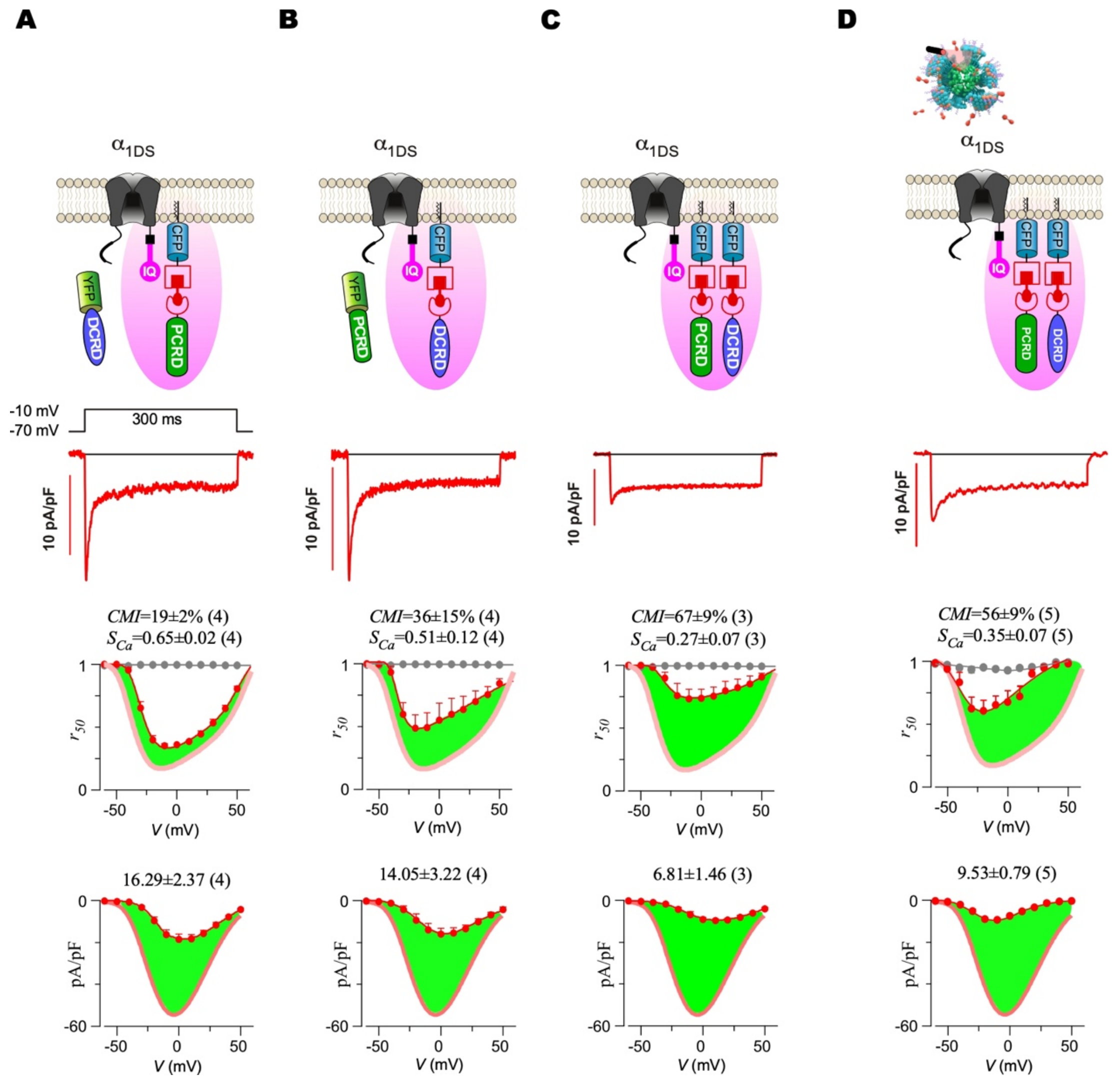


**Figure S7. End-point evaluations of membrane-assisted CMI by P+D peptides.**

(A-C) Endpoint profiles of $\alpha_{1DS}$ currents for chemogenetic CMI. The design schemes of A-C are identical to Figure S5A, S5D and Figure 5B, respectively. Cells were treated with bath solution containing 1 μM rapamycin for at least 10 min, then recorded with the voltage protocol. The representative $I_{Ca}$ traces, inactivation ($r_{50}$) and activation ($I_{peak}$, pA/pF) profiles across the full range of $V$ are shown. The values of $S_{Ca}$, $CMI$ and $I_{peak}$ are indicated in the plots as the indices of CDI, CMI and activation. The curves in pale red represent the $r_{50}$ and $I_{peak}$ profiles of $\alpha_{1DS}$. Green areas indicate the differences from $\alpha_{1DS}$.

(D) Endpoint profiles of $\alpha_{1DS}$ currents for opto-chemogenetic CMI. The design scheme is identical to Figure 5D. Cells are treated with bath solution containing rapamycin@PDPP, and irradiated by 808 nm laser *in situ* for 5 min to release the rapamycin with a final concentration of 1 μM. All statistical data are given as mean ± SEM.

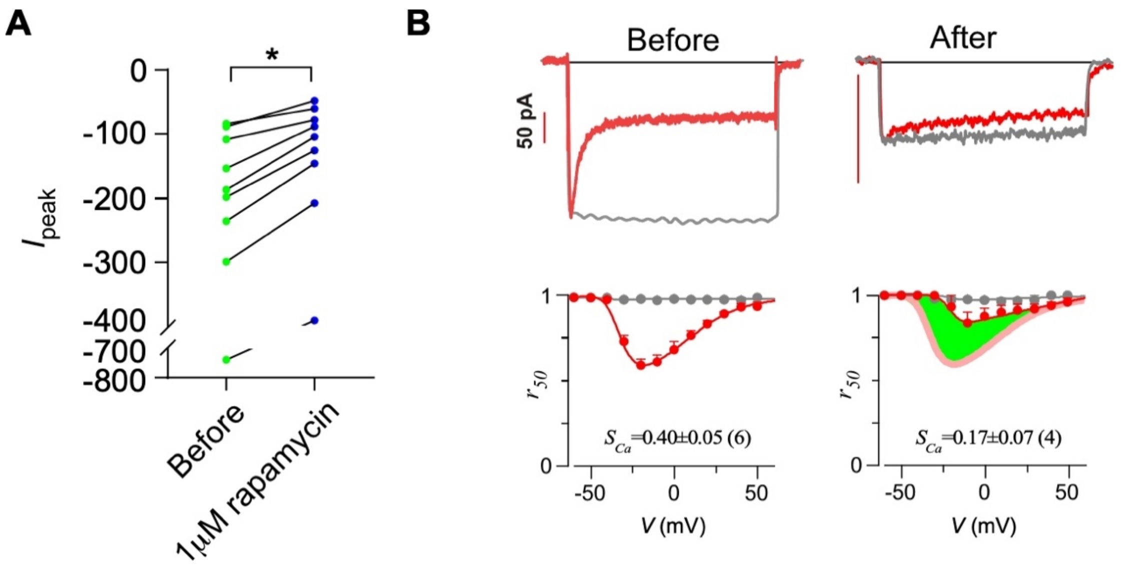


**Figure S8. Inducible membrane-assisted CMI on $\alpha_{1DL}$ by P-D peptides.**

(A) Peak currents ($I_{peak}$, pA) of $\alpha_{1DL}$ at the membrane potential ($V$) of -10 mV before and 300 s after 1 μM rapamycin, in accordance with Figure 6A-6C.

(B) FRB-CFP-Ras and FKBP-YFP-$P_CD_C$ (P-D) were co-expressed with $\alpha_{1DL}$. Representative current traces (upper) and inactivation $r_{50}$ profiles (lower) for the control group (left) or the group of 1 μM rapamycin (right).

All statistical data are given as mean ± SEM. Paired $t$-test was applied for $A$: $*p<0.05$.

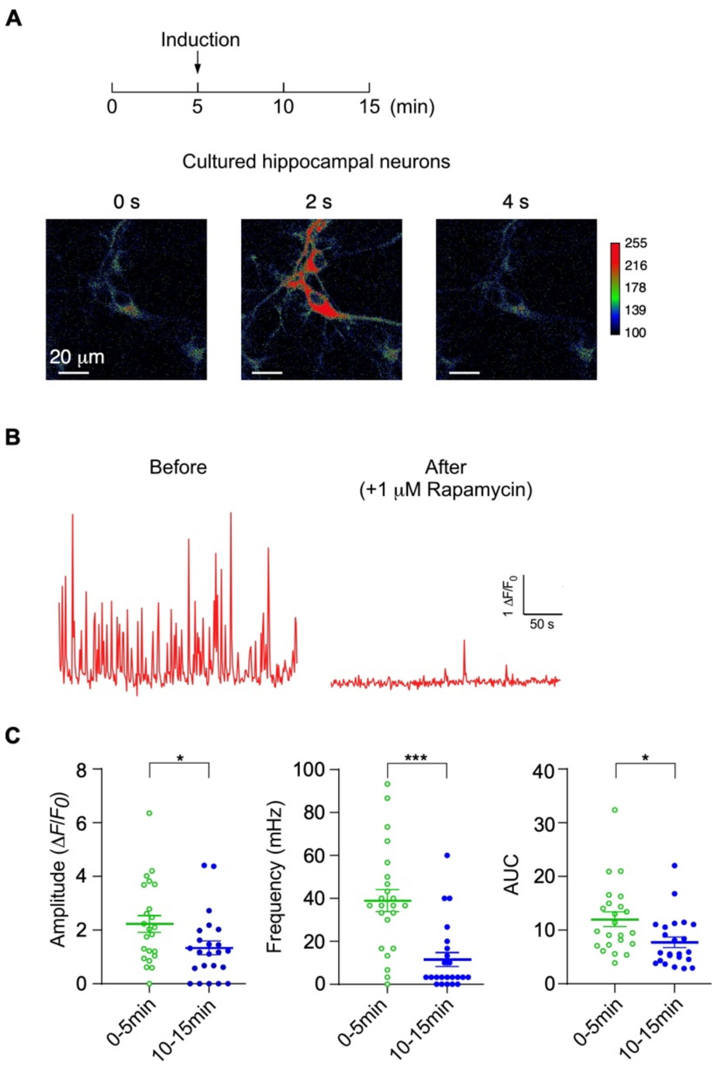


**Figure S9. Inducible P-D peptides suppressed oscillatory $Ca^{2+}$ in hippocampal neurons.** (A) Experimental protocol for acute inhibition on cultured hippocampus neurons from newborn rats. 1 μM rapamycin (indicated by arrow) was applied to the hippocampal neurons expressing membrane-anchored FRB-CFP-Ras and cytosolic FKBP-YFP-$P_CD_C$ at DIV 7. Confocal imaging was performed before and after rapamycin treatment at the indicated timepoints. Exemplary images, color-coded to indicate fluorescence intensity, illustrate $Ca^{2+}$ dynamics.

(B) Spontaneous $Ca^{2+}$ activities before (left) or after applying 1 μM rapamycin (right).

(C) Summary for $Ca^{2+}$ dynamics in neurons before (in green) or after 1 μM rapamycin (in blue), respectively, with the key indices including the average frequency (mHz, left), peak amplitude ($\Delta F/F_0$, middle), and AUC ($\Delta F/F_0$ per min, right).

All statistical data are given as mean ± SEM. Two-tailed Student's *t*-test was applied: *, $p$<0.05; ***, $p$<0.001.

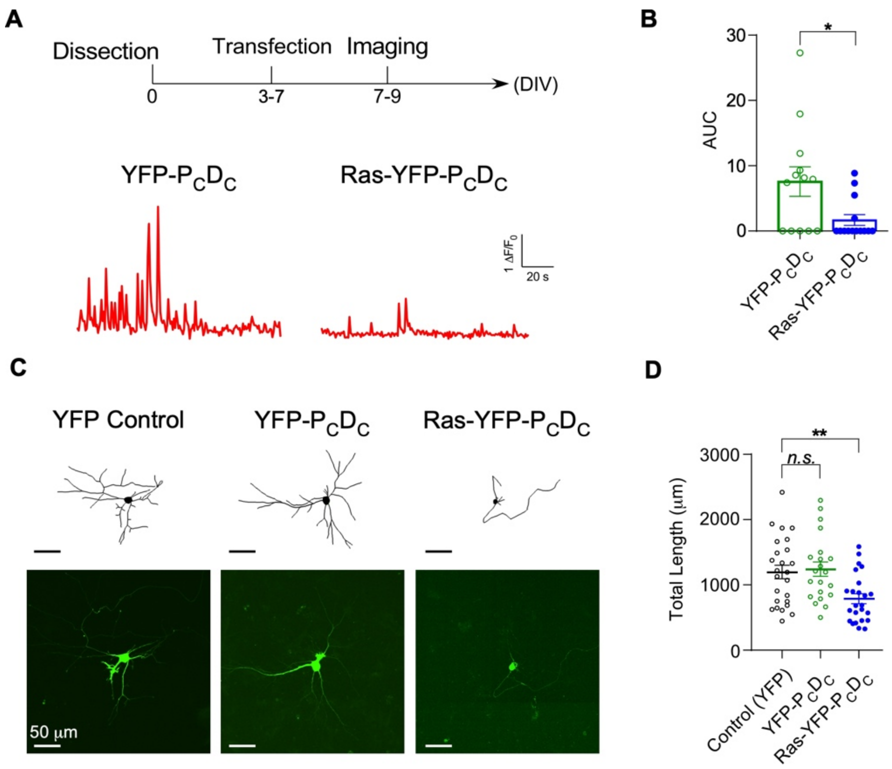


**Figure S10. Membrane-anchored P-D peptides inhibited $Ca_V1$ influx- neuritogenesis coupling in hippocampus neurons.**

(A) Cultured hippocampus neurons were transiently transfected on DIV 3-7 with YFP, FKBP-YFP-$P_CD_C$, or Ras-FKBP-YFP-$P_CD_C$, respectively. Confocal microscopy was conducted at DIV 7-9. Representative $Ca^{2+}$ dynamics for neurons expressing FKBP-YFP-$P_CD_C$ or Ras-FKBP-YFP-$P_CD_C$.

(B) AUC ($\Delta F/F_0$ per min) analysis for hippocampus neurons expressing FKBP-YFP-$P_CD_C$, or Ras-FKBP-YFP-$P_CD_C$, respectively.

(C, D) Neurite tracing, fluorescence imaging (C) and total length analysis (D) for hippocampus neurons expressing YFP, FKBP-YFP-$P_CD_C$, or Ras-FKBP-YFP-$P_CD_C$, respectively.

All statistical data are given as mean ± SEM. The significance tests include two-tailed Student's *t*-test (B) and one-way ANOVA followed by Dunnett for post hoc test (D): *, $p<0.05$; **, $p<0.01$; n.s., $p>0.05$.
